# Crystal structure of the M_5_ muscarinic acetylcholine receptor

**DOI:** 10.1101/730622

**Authors:** Ziva Vuckovic, Patrick R. Gentry, Alice E. Berizzi, Kunio Hirata, Swapna Varghese, Geoff Thompson, Emma T. van der Westhuizen, Wessel A.C. Burger, Raphaёl Rahmani, Celine Valant, Christopher J. Langmead, Craig W. Lindsley, Jonathan Baell, Andrew B. Tobin, Patrick M. Sexton, Arthur Christopoulos, David M. Thal

## Abstract

The human M_5_ muscarinic acetylcholine receptor (mAChR) has recently emerged as an exciting therapeutic target for treating a range of disorders, including drug addiction. However, a lack of structural information for this receptor subtype has limited further drug development and validation. Here we report a high-resolution crystal structure of the human M_5_ mAChR bound to the clinically used inverse agonist, tiotropium. This structure allowed for a comparison across all five mAChR family members that revealed important differences in both orthosteric and allosteric sites that could inform the rational design of selective ligands. These structural studies together with chimeric swaps between the extracellular regions of the M_2_ and M_5_ mAChR further revealed the structural insight into “kinetic-selectivity”, where ligands show differential residency times between related family members. Collectively, our study provides important insights into the nature of orthosteric and allosteric ligand interaction across the mAChR family that could be exploited for the design of selective ligands.

**Significance Statement:** The five subtypes of the muscarinic acetylcholine receptors (mAChRs) are expressed throughout the central and peripheral nervous system where they play a vital role in physiology and pathologies. Recently, the M_5_ mAChR subtype has emerged as an exciting drug target for the treatment of drug addiction. We have determined the atomic structure of the M_5_ mAChR bound to the clinically used inverse agonist tiotropium. The M_5_ mAChR structure now allows for a full comparison of all five mAChR subtypes and reveals subtle differences in the extracellular loop (ECL) regions of the receptor that mediate orthosteric and allosteric ligand selectivity. Together these findings open the door for future structure-based design of selective drugs that target this therapeutically important class of receptors.

## Introduction

The muscarinic acetylcholine (ACh) receptors (mAChRs) are Class A G protein-coupled receptors (GPCRs) that together with the nicotinic acetylcholine receptors facilitate the actions of the neurotransmitter, ACh, throughout the body. The mAChR family comprises five subtypes where M_1_, M_3_, and M_5_ are preferentially coupled to the G_q/11_ protein-mediated signalling pathways, and M_2_ and M_4_, show preference for G_i/o_ protein-dependent signalling. Localization studies have revealed that the mAChR subtypes are differentially distributed, with M_1_, M_4_, and M_5_ mAChRs found predominantly in the central nervous system (CNS), where they are essential for normal neuronal function, while M_2_ and M_3_ mAChRs are expressed more widely, including in the periphery, where they are involved in cardiovascular as well as gut motility and secretory processes (1).

Given the involvement of mAChRs in such a wide range of fundamental physiological processes, they have long been valued as targets for novel therapeutics, in particular the central M_1_ and M_4_ mAChRs, which have garnered attention due to their involvement in cognition and memory (2). In contrast, relatively less is known about the M_5_ mAChR subtype, which represents less than 2% of the total CNS mAChR population (3, 4). Despite its low level of expression, this receptor plays a vital role in the mesolimbic reward pathway due to its presence on dopaminergic neurons of the ventral tegmental area (VTA) (5–8). Additionally, there is a large population of non-neuronal M_5_ mAChRs located within the endothelium of the cerebral vasculature, suggesting that the receptor may modulate cerebral vasodilatory processes (9, 10). These observations correlate well with phenotypic data from M_5_ mAChR knockout mice where the cerebral vasculature is constitutively constricted, resulting in decreased cerebral blood flow (11, 12). Additionally, M_5_ mAChR knockout mice exhibited attenuated reward-seeking behaviour to drugs of addiction, such as cocaine and morphine in self-administration and conditioned place-preference experiments (13–15). Moreover, in recent studies involving rats (16–18), ethanol-seeking behaviour and oxycodone self-administration were attenuated by the selective M_5_ mAChR negative allosteric modulator (NAM) ML375 (19). From these studies, the M_5_ mAChR has emerged as a potential target for the treatment of drug addiction.

Despite such promising data, further study of the M_5_ mAChR has been hindered by a lack of selective small molecule tool compounds. Designing conventional small molecule ligands that target the orthosteric ACh binding site of individual mAChR subtypes has been challenging due to the highly conserved sequence homology of the mAChR orthosteric site residues (1), and in part due to a lack of detailed structural information for all five receptor subtypes. While structures of the M_1_–M_4_ mAChRs have been previously determined, there are no available structures for the M_5_ mAChR. Therefore, to provide a complete structural comparison of all five family members and to gain insight into ligand binding we determined a high-resolution crystal structure of the M_5_ mAChR.

## Results

### Crystallization and determination of the M_5_ mAChR structure

To determine the M_5_ mAChR structure we designed a construct where residues 225-430 of intracellular loop 3 were removed and replaced with a T4 lysozyme (T4L) fusion protein. Additionally, to promote crystallization, the first 20 N-terminal amino acids were cleaved by a Tobacco Etch Virus (TEV) protease site engineered into the receptor (Supplementary Figure 1a). The inverse agonist, tiotropium, was used to stabilize the inactive state as it has a slow dissociation rate at the M_5_ mAChR (20), and was also used in the determination of the M_1_, M_3_, and M_4_ mAChR structures (21, 22). The M_5_-T4L•tiotropium complex was crystallized in lipidic cubic phase (LCP), and crystals were obtained within 1-2 days; however, despite many rounds of optimization, diffraction was limited to 7 Å. To improve the resolution we built upon a study from Yasuda *et al*. (23) that predicted that mutation of the amino acid at position 3.39 (numbered according to Ballesteros-Weinstein (24)) to Arg would create a thermostabilized receptor by promoting an ionic bond between this residue and the highly conserved D^2.50^ residue. Recently, the same S^3.39^R mutation was applied to the M_2_ mAChR resulting in a series of higher resolution structures (25). Although introduction of the S117^3.39^R mutation resulted in a construct that binds the antagonists NMS or tiotropium with a slightly reduced affinity relative to the WT M_5_ mAChR, the effect of the mutation on reducing ACh affinity was substantially more pronounced (Supplementary Figure 1b-e), consistent with the ability of the construct to favour an inactive over an active state. Similar differential effects on antagonist versus agonist affinity were previously observed for S^3.39^R at the M_2_ mAChR (25). Notably, introduction of the S117^3.39^R mutation increased our M_5_ mAChR yields during purification and resulted in crystals that diffracted to a resolution of 3.4 Å. Data were collected from approximately 130 crystals, and the structure was determined by molecular replacement using the M_3_ structure (PDB: 4U15) and an ensemble of T4L structures as templates (Figure 1a, Supplementary Table 1).

**Figure 1.**
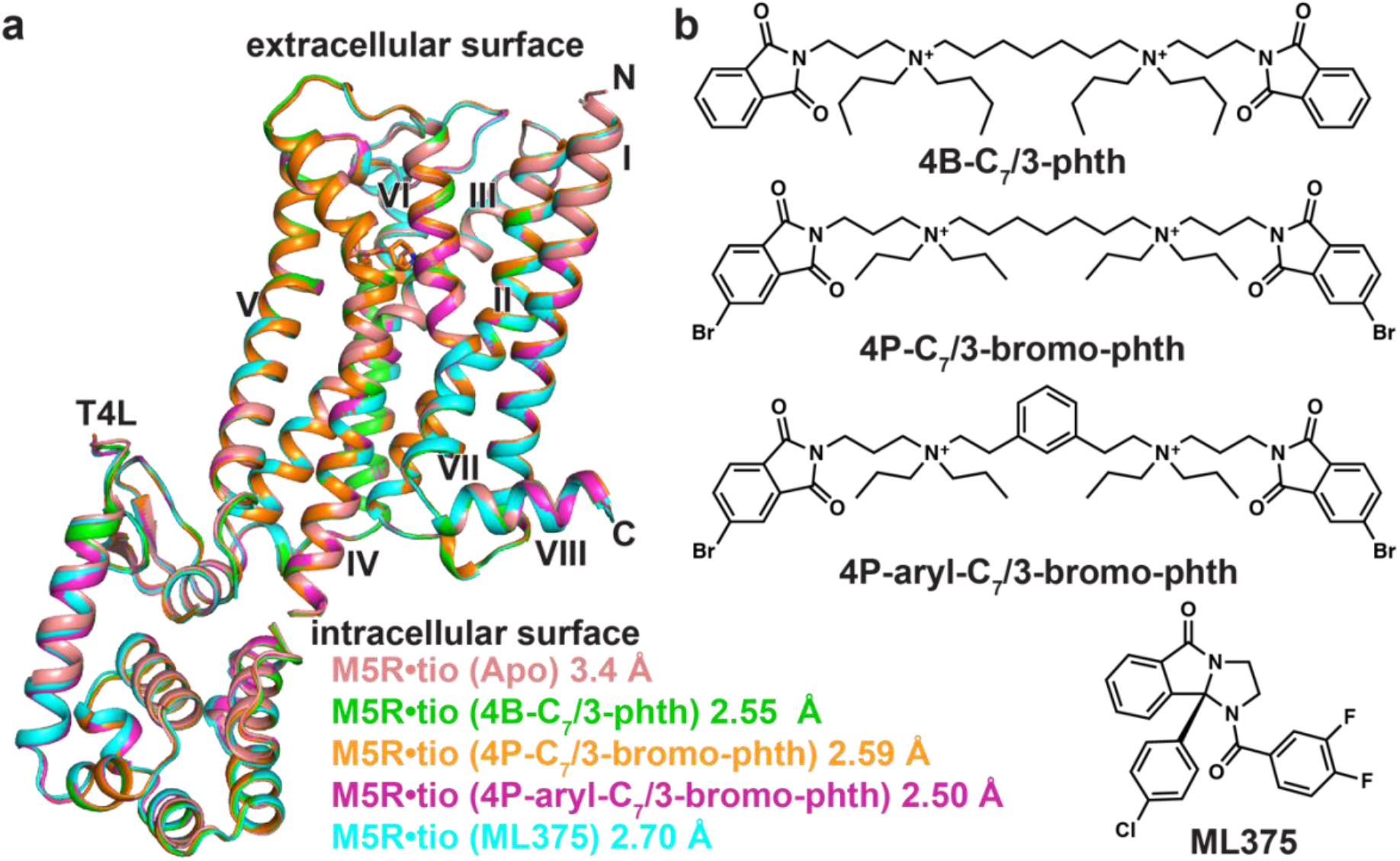
Structures of M_5_-T4L bound to tiotropium. **(a)** Overlay of five different M_5_ mAChR structures determined in the presence of tiotropium and **(b)** different allosteric modulators. The structure from 4B-C_7_/3-phth was the most resolved of all the datasets and is used in all further comparisons (Supplementary Table 1).

To investigate the nature of NAMs binding to the M_5_ mAChR, we attempted to obtain co-crystal structures. Given that the bis-ammonium alkane type ligands tend to have higher affinities for the M_5_ mAChR than the prototypical modulator, gallamine (26), we tried to obtain a ternary complex structure of the M_5_ mAChR with tiotropium and several bis-ammonium alkane ligands (Figure 1b). We initially used the modulator, 4B-C_7_/3-phth, which resulted in crystals that grew to a much larger size and diffracted to a resolution of 2.55 Å (Figure 1, Supplementary Table 1, Supplementary Figure 2). Based on previous data (27), we predicted that 4B-C_7_/3-phth would bind in the extracellular vestibule (ECV). While there were regions of strong electron density present in the ECV, we could not unambiguously model 4B-C_7_/3-phth into the density as a molecule of the precipitant, polyethylene glycol 400 (PEG400), also likely binds in this site (22, 28), and may explain why researchers have had difficulty in obtaining co-NAM bound structures for the mAChRs.

Subsequently, we designed two new bis-ammonium alkane analogs using the higher affinity 4P-C_7_/3-phth scaffold (27) to try and improve modulator affinity (Figure 1b, Supplementary Figure 3a, Supplementary Table 2) and detectability by X-rays. The first modification added two bromine atoms (4P-C_7_/3-bromo-phth) to increase the size of the pthalamide groups (29), and the second modification rigidified the flexible 7-carbon linker with an aromatic hydrocarbon (4P-aryl-C_7_/3-bromo-phth). Both ligands displayed an increased affinity for the M_5_ mAChR versus 4B-C_7_/3-phth, but had a slightly reduced affinity in relation to the parent compound (4P-C_7_/3-phth) when assayed for NAM activity in inhibiting [^3^H]NMS radioligand binding (Supplementary Figure 3a, Supplementary Table 2). Like 4B-C_7_/3-phth, the addition of either 4P-C_7_/3-bromo-phth or 4P-aryl-C_7_/3-bromo-phth to purified M_5_ mAChR and reconstitution into LCP yielded crystals that diffracted to a higher resolution (Supplementary Table 1). A full data set for 4P-aryl-C_7_/3-bromo-phth was collected at wavelength of 0.92 Å to maximize the anomalous Br signal in a single wavelength anomalous diffraction experiment, however, no such signal was detected, suggesting that 4P-aryl-C_7_/3-bromo-phth was not present in the structure. Since the structure was solved by merging a large number of datasets, there is possibility that the Br signal for the NAM would be averaged out if NAM occupancy is low. However, inspection of different datasets did not indicate that this was the case.

As an alternate strategy, we attempted to determine a co-crystal structure with the structurally diverse M_5_ mAChR selective NAM, ML375 (19). In comparison to the bis-ammonium ligands, the addition of ML375 resulted in a slightly lower resolution structure (2.7 Å, Supplementary Table 1) and, as was the case with the bis-ammonium NAMs, we were not able to assign ML375 into any electron density. Comparison of all M_5_ mAChR structures showed that they were nearly identical, with root mean square deviation values of 0.09–0.22 Å. The higher resolution 2.55 Å M_5_•tiotropium (4B-C_7_/3-phth) structure was used for further comparison, as this was the best resolved and modelled structure.

### Family-wide comparison of all mAChR subtypes

The solution of the M_5_ mAChR structure allows the first complete subtype-wide comparison of this important GPCR family. The structure of the M_5_ mAChR is similar to the previously determined structures of the M_1_-M_4_ mAChR subtypes (21, 22, 30) with a root mean squared deviation of 0.5-0.8 Å (Figure 2a) for the seven-transmembrane domain across all subtypes. The five mAChR subtypes are most similar in the orthosteric binding site, which is the most conserved region of the receptor. The fact that our M_5_ mAChR structure was obtained in complex with the same ligand (tiotropium), as the M_1_, M_3_ and M_4_ mAChR structures, allowed for specific, detailed comparison of residues lining this orthosteric binding site (Figures 2b,c).

**Figure 2.**
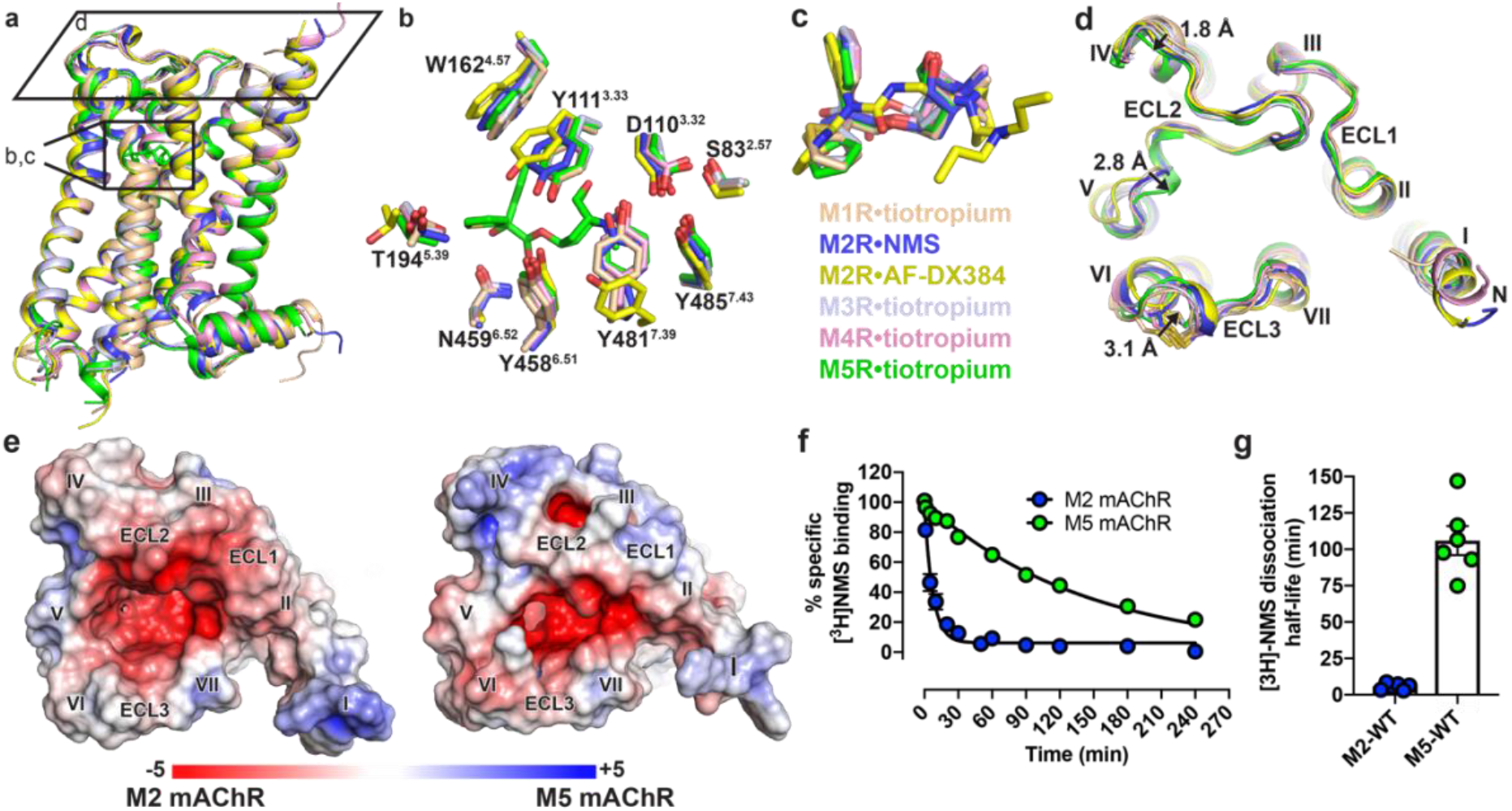
Structural comparison of M_1_-M_5_ mAChRs. **(a)** The overall view of the M_1_-M_5_ mAChR structures aligned to the M_5_ mAChR and shown as cartoons. M_1_•tiotropium is coloured peach (PDB: 5CXV), M_2_•NMS dark blue (5ZKC), M_2_•AF-DX384 yellow (5ZKB), M_3_•tiotropium light blue (4U15), M_4_•tiotropium pink (5DSG) and M_5_•tiotropium green (6OL9). **(b)** Comparison of residues (stick representation) lining the orthosteric site with tiotropium from the M_5_ mAChR displayed, and **(c)** overlay of the orthosteric ligands. **(d)** View from the extracellular surface comparing differences in the ECL regions across the M_1_-M_5_ mAChRs. Distances between the backbone of M_1_ and M_5_ mAChR residues in ECL2 and ECL3 are shown and indicated by arrows. **(e)** Electrostatic and surface potential of M_2_ and M_5_ mAChR (+5kT/e in blue and −5kT/e in red) mapped on the surface of the receptors calculated at pH 7.0 using PDB2PQR and APBS (31). **(f)** Comparison of dissociation rate and **(g)** dissociation half-life of [^3^H]NMS by the addition of 10 μM atropine at the M_2_ and M_5_ mAChRs.

This comparison demonstrated that the residues within the orthosteric pocket are absolutely conserved between the receptors. Although there is no tiotropium bound M_2_ mAChR structure, there are now six different inactive state M_2_ mAChR structures, which include structures bound with the non-selective ligands QNB and NMS, and the M_2_ mAChR selective ligand, AF-DX384 (25). The 2.3 Å M_2_•NMS structure is most similar to the tiotropium bound mAChR structures, though residues Y^3.33^ and Y^7.39^ of the “tyrosine lid” (Y^3.33^, Y^6.51^, and Y^7.39^) are positioned in a distinct conformation in comparison to the tiotropium bound structures. These differences in the tyrosine lid positions are more pronounced in the M_2_•AF-DX384 structures, allowing the accommodation of this bulkier ligand into the orthosteric binding pocket (Figures 2b-c).

Subtle yet notable differences between the mAChR subtypes are observed for ECL2 and ECL3, corresponding to regions that are the least conserved across the receptors (Figure 2d). At ECL2 there is a 1.8 Å difference across all five subtypes beginning at the first non-conserved residue of ECL2. As ECL2 progresses towards TM5, a conserved 3_10_ helix motif moves inward by 2.8 Å in the M_5_ mAChR when compared to the M_1_ structure. Similarly, the conserved ECL3 disulphide bond is displaced inwards by 3.1 Å for the M_3_ and M_5_ mAChRs (Figure 2d), relative to the other subtypes. These observed differences in the positons of ECL2 and ECL3, along with differences in amino acid composition contribute to a more constricted entrance to the orthosteric binding site at the M_5_ (and M_3_) versus the M_2_ mAChR (Figure 2e). Furthermore, this contraction of the entrance in the antagonist-bound structures may contribute to the slower dissociation rate of orthosteric ligands from the M_5_ and M_3_ mAChRs, in comparison to other subtypes like the M_2_ mAChR. For example, despite having similar equilibrium binding affinities, [^3^H]NMS dissociates 18-fold more slowly at the M_5_ than the M_2_ mAChR with half-lives of dissociation of 100 ± 11.6 and 5.6 ± 1.2 min, respectively (Figure 2f,g).

### Structural differences between the extracellular vestibules (ECVs) of the M_2_ and M_5_ mAChRs

An alternative strategy to generating selective ligands is to target non-conserved allosteric sites (32). This has been extensively explored for the mAChR family where a palette of both positive and negative allosteric modulators has been identified (33, 34). Structural and mutagenesis studies have established that many of these ligands bind to a “common” allosteric site that is located above the orthosteric site and within an ECV (Figure 3, Supplementary Figure 4) (35). In fact, the M_5_ mAChR has often served as model system for early research into understanding the binding mode and mechanism of selectivity for prototypical modulators, such as the bis-ammonium alkane ligands (Figure 1b), that have higher sensitivity for modulating the M_2_ mAChR and lower sensitivity for the M_5_ mAChR (26, 36–39). These studies identified non-conserved residues in ECL2 (P179^−4^, E182^−1^, and Q184^+1^; superscript indicates the position of ECL2 residues relative to the conserved Cys in ECL2) and TM7 (V474^7.32^ and H484^7.36^) as residues that can account for M_2_/M_5_ subtype selectivity. Comparison of the ECV between the M_2_ and M_5_ mAChRs confirm differences in the orientations and positions of these residues that could mediate the selectivity. Namely, P179^−4^ in ECL2 restricts the position of E182^−1^ forcing the residue into the ECV near Q184^+1^. Residue Q184^+1^, which is a F/Y residue for the M_1_-M_4_ mAChRs subtypes, is a key residue for the activity of many allosteric modulators. Other major differences between the M_2_/M_5_ ECVs are in the positions of non-conserved residues lining the top of TM6 starting from S465^6.58^ across ECL3 and down to residue H478^7.36^ in TM7. At the M_5_ mAChR these residues are bulkier and point more inward constricting the overall size of the ECV (Figure 3).

**Figure 3.**
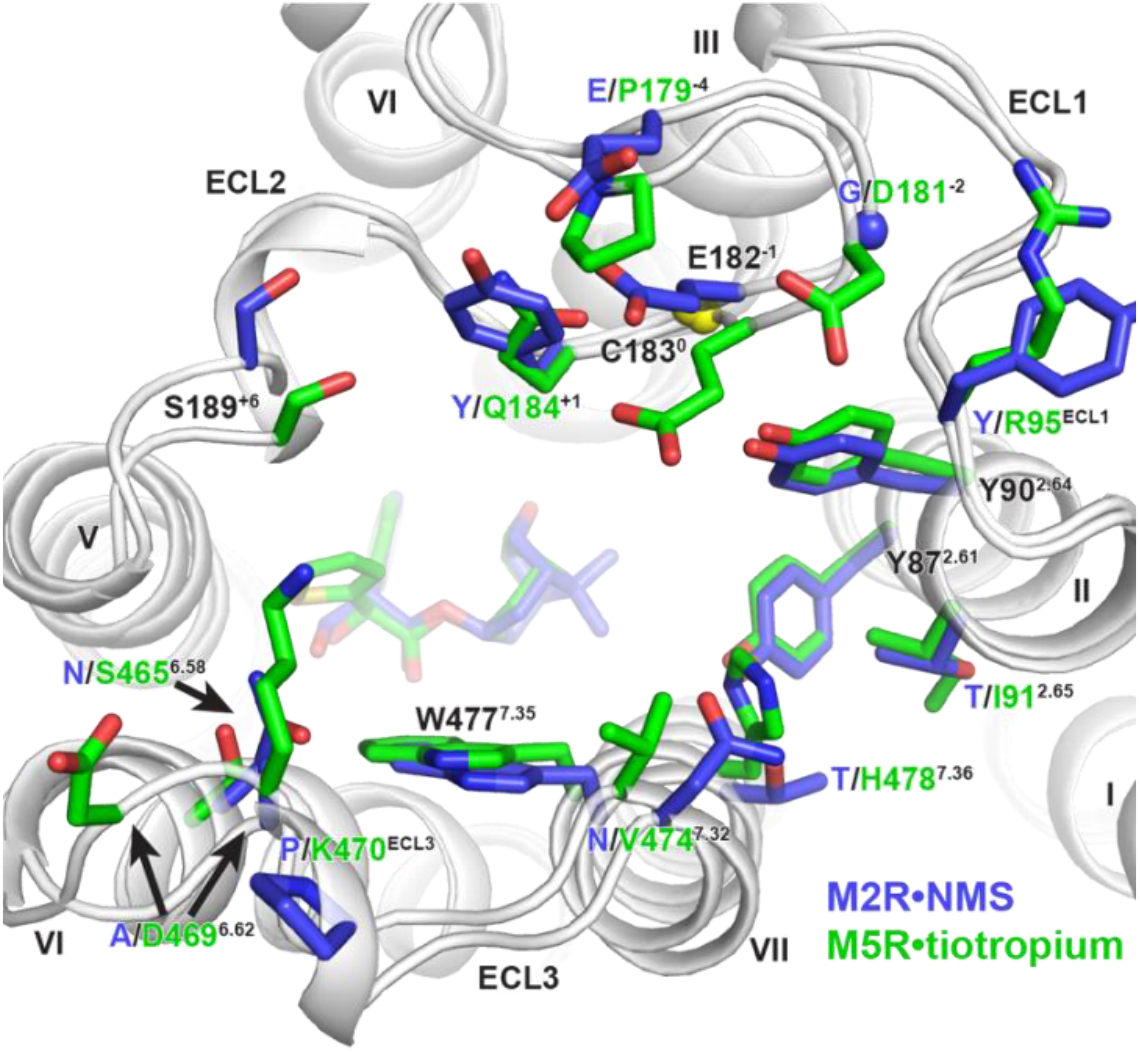
Comparison of residues lining the extracellular vestibule of the M_2_ and M_5_ mAChR. M_2_•NMS is shown in dark blue and M_5_•tiotropium in green. Conserved residues are labelled black and non-conserved residues are coloured according to receptor subtype. Residues are numbered based on the M_5_ mAChR, with residues in ECL2 numbered relative to the conserved cysteine in ECL2, which is shown as a yellow sphere.

### Role of the M_5_ and M_2_ mAChR ECL regions in orthosteric and allosteric ligand binding

The effect of ECL regions on orthosteric ligand access and egress has significant biological and clinical relevance (40). Therefore, to investigate the role of the ECLs on modulating the slower dissociation kinetics of the M_5_ mAChR in comparison to the M_2_ mAChR, we designed full ECL1, ECL2, and/or ECL3 chimeric swaps between the two subtypes (Figure 4). The ECL chimeras had similar levels of expression and binding of [^3^H]NMS to wild type receptors (Supplementary Table 3). As previously noted, the M_2_ mAChR has a shorter half-life for [^3^H]NMS dissociation in comparison to the M_5_ mAChR (Figure 2f). Incorporation of the M_2_ ECL1 or ECL3 into the M_5_ mAChR increased [^3^H]NMS dissociation, while the reciprocal chimeric swap decreased [^3^H]NMS dissociation at the M_2_ mAChR. Unexpectedly, it was the ECL1 swaps that had the largest effect on [^3^H]NMS dissociation between the two subtypes, particularly at the M_5_ mAChR (Figure 4, Supplementary Table 4). A possible structural explanation for this observation could be that R95^ECL1^, which is a conserved Tyr residue at the M_1_-M_4_ subtypes, is capable of forming an ionic bond with either the M_5_ ECL2 residue, D181^−2^, or in the case of the M_2_ ECL1 chimera residue D173^−3^ (Figure 3, Supplementary Figure 4). Such an interaction could tether ECL1 and ECL2 limiting their overall dynamics and thus reduce rates of orthosteric ligand dissociation. It is important to note that R95^ECL1^ is involved in an ionic interaction mediated through the crystal lattice with a neighbouring T4L molecule (Supplementary Figure 2d-f), and as a result it does not directly interact with D181^−2^ in the M_5_ mAChR structure though it is well positioned to do so.

**Figure 4.**
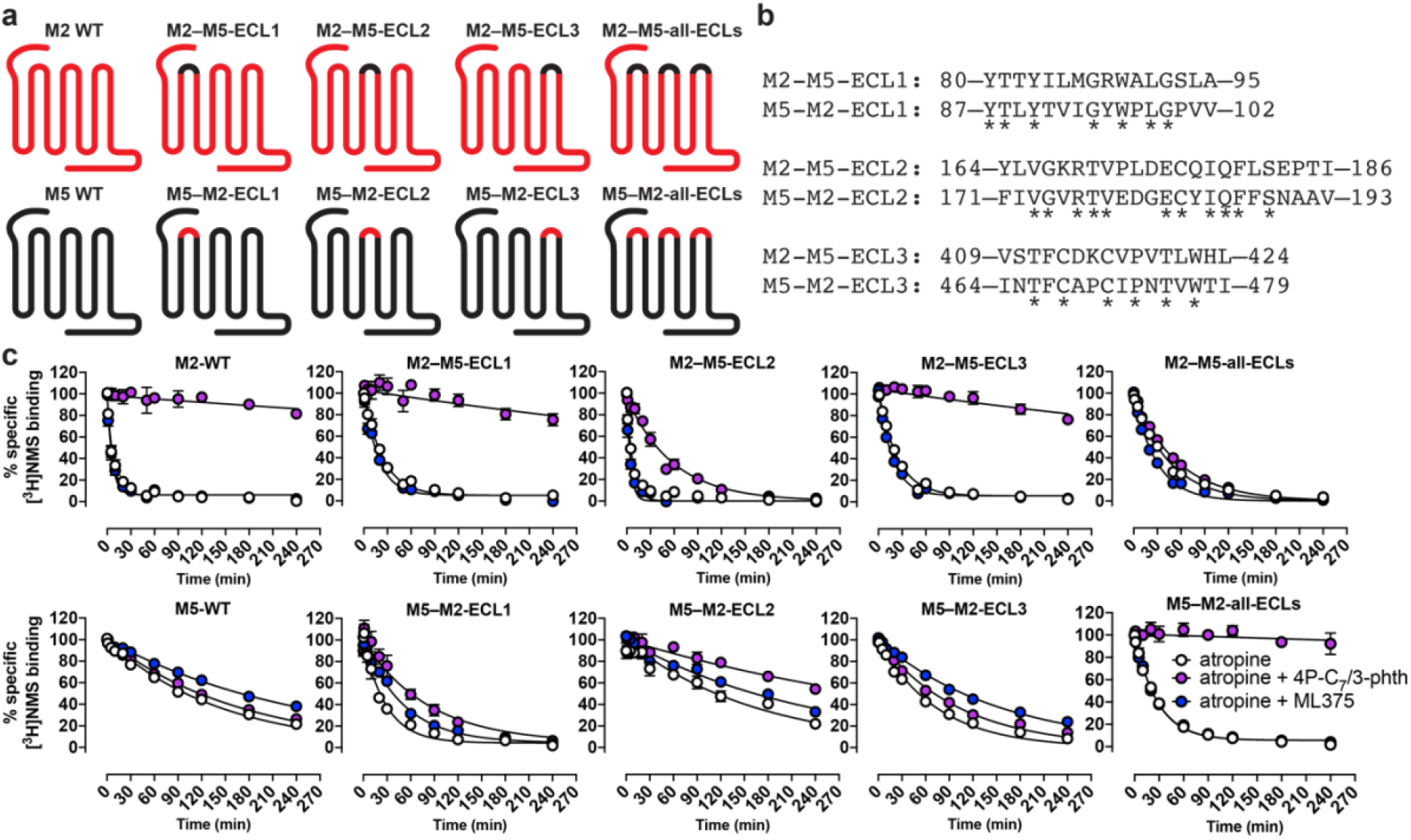
[^3^H]NMS binding dissociation kinetic studies of chimeric swaps between the ECLs of the M_2_ and M_5_ mAChRs. **(a)** Cartoons and **(b)** amino acid sequence composition for the M_2_ and M_5_ ECL chimeras used in this study, with conserved residues noted by an asterix. **(c)** [^3^H]NMS re-association was prevented by the addition of 10 μM atropine, and radioligand dissociation was monitored in the absence or presence of 10 μM ML375 or 10 μM 4P-C_7_/3-phth. Data points represent the mean ± S.E.M. of three or more independent experiments performed in duplicate. Quantitative parameters derived from this experiment are listed in Supplementary Table 4.

A hallmark feature of an allosteric ligand that modulates orthosteric ligand affinity is the ability to either increase or decrease the rate of dissociation of an orthosteric ligand. To examine the effect of allosteric modulators on NMS dissociation across the M_5_ and M_2_ ECL chimeras, we used the bis-ammonium alkane ligand 4P-C_7_/3-phth, which had been previously studied at the M_2_ mAChR and had high affinity for the M_5_ mAChR (Supplementary Table 2), or the M_5_ selective modulator ML375 (19, 27). In the presence of ML375, [^3^H]NMS dissociation was reduced at the M_5_ mAChR and had no effect at the M_2_ mAChR, whereas the addition of 4P-C_7_/3-phth reduced radioligand dissociation at the M_2_ mAChR but not at the M_5_ mAChR (Figure 4, Supplementary Table 4). The ECL1 and ECL3 chimeric swaps had little effect on the activity of ML375 for either receptor subtype, and slightly increased the activity of 4P-C_7_/3-phth at the M_5_ mAChR. For the ECL2 chimeras, there was no effect on activity of ML375. However, there was a loss of 4P-C_7_/3-phth activity at the M_2_ mAChR and a corresponding gain of activity at the M_5_ mAChR. These results are in line with previous studies and highlight the importance of residues in ECL2, particularly M_2_-Y177 and M_5_-E184, on modulating the activity of bis-ammonium alkane ligands. Interestingly, when all three ECLs were swapped, the resulting M_2_ and M_5_ chimeric constructs functioned more like their swapped receptor counterpart. That is, for the M_2_–M_5_-all-ECL construct, 4P-C_7_/3-phth had little effect, and although ML375 did not retard [^3^H]NMS dissociation, it slightly increased the rate of [^3^H]NMS dissociation suggesting an allosteric mode of action (Figure 4, Supplementary Table 4). Conversely, for the M_5_–M_2_-all-ECL construct, 4P-C_7_/3-phth retarded radioligand dissociation and, surprisingly, ML375 had no effect. While none of the chimeric constructs ever fully switched the basal dissociation rate of [^3^H]NMS or ML375 activity to that observed for the corresponding WT constructs, the data nonetheless suggest that the ECL regions modulate the overall conformation of mAChRs and directly influence the dissociation of ligands from the orthosteric site.

## Discussion

Individual mAChR subtypes have long been pursued as drug targets for a range of CNS disorders, and recent studies have begun to validate the M_5_ mAChR as a novel target for the treatment of drug addiction (4, 41). In this study, we have determined a high-resolution crystal structure of the M_5_ mAChR, thus allowing the first subtype-wide comparison for any aminergic GPCR subfamily. Introduction of the inactive state stabilizing mutation S117^3.39^R, which was recently used to stabilize the M_2_ mAChR (25), was crucial to obtaining well-diffracting crystals and suggests that this mutation could be applied to aid the determination of inactive state structures for other related GPCRs. We further improved the resolution of the M_5_ mAChR structure by adding allosteric modulators to the purified protein prior to crystallization. Despite the consistent increase in resolution that each of the allosteric modulator provided, we were not able to model any of the modulators into electron density. From a pharmacological perspective, a lack of modulator binding is not surprising, as all of the modulators tested in this study showed strong negative cooperativity with tiotropium (Supplementary Figure 3b). Nevertheless, it is still paradoxical that the addition of an allosteric modulator can clearly improve receptor crystallization and diffraction, though not be visible in any resulting structures. This phenomenon has been noted at other GPCRs, such as the M_2_ mAChR that was crystallized in the presence of the modulator, alcuronium, and the CC chemokine receptor 2A that was crystallized in the presence of the modulator, AZD-6942, but where neither modulator could be observed in the resulting structures (25, 42).

Comparison of all five mAChR structures further confirms the well conserved transmembrane core and orthosteric binding site that has made the discovery of highly selective drugs for these receptor subtypes incredibly challenging. The most apparent structural differences between the mAChR subtypes is in the ECL regions. Though these differences are generally quite subtle they are important because they open up the possibility for designing selective molecules in a way that has not previously been possible (Supplementary Figure 4). For example, a recent crystal structure of the M_2_ mAChR bound to the M_2_-selective antagonist AF-DX384 revealed that selectivity is mediated by differential interactions between the ligand and residues in ECL2, which lead to an outward displacement in ECL2 and the top of TM5 (Figure 2d) (25). Likewise, by utilizing knowledge of a single amino acid difference in ECL2 between the M_2_ and M_3_ mAChRs, molecular docking and structure-based design led to the discovery of a new M_3_-selective antagonist with 100-fold selectivity over the M_2_ mAChR (43). These results are similar to the structure-based design of biased ligands targeting the D2 dopamine receptor that were designed by utilizing specific amino acid-ligand contacts in ECL2 and TM5 (44). Taken together, these findings indicate that the differential targeting of ECL residues may be a path forward for creating selective mAChR ligands. This is well supported by the fact that many mAChR-selective allosteric modulators interact with the ECL regions (27, 35), and suggest that designing orthosteric ligands linked to allosteric pharmacophores, known as bitopic ligands, is a potential strategy for future structure-based drug design.

Drug discovery has typically focused on optimizing ligand affinity and selectivity. However, it is now apparent that binding kinetics are just as important (40, 45–47). This is illustrated two ways with the drug tiotropium as a pertinent example. First, tiotropium has slow rate of dissociation from the M_3_ mAChR, which is a key feature of the drug that allows for a once daily dosing for the treatment of chronic obstructive pulmonary disease (48). Second, though tiotropium has the same equilibrium binding affinity for the M_3_ and M_2_ mAChRs, it exhibits kinetic selectivity for the M_3_ over M_2_ mAChR, by having substantially different rates of dissociation. This kinetic selectivity over the M_2_ mAChR is postulated to be due to differences in the electrostatics and dynamics of the ECL region (48). The M_5_ mAChR is similar to the M_3_ mAChR with respect to having slow rates of orthosteric ligand dissociation (20), and data from our M_2_/M_5_ ECL chimeras support the idea of the ECL regions mediating kinetic selectivity as [^3^H]NMS dissociation was switched between the M_2_ and M_5_ mAChRs (Figure 4). Notably, none of the combined ECL chimeras could ever fully switch the dissociation kinetics between subtypes, suggesting that other mechanisms are operative such as the global conformation of the ECLs. Our results also highlight the importance of the ECL regions on conferring sensitivity to allosteric modulators across different subtypes. By swapping out the entire ECL region between the M_2_ and M_5_ mAChRs we were able to completely alter the sensitivity of a modulator that is selective for the M_2_ versus the M_5_ mAChR and vice versa. These results support the importance of the ECL region for mediating ligand selectivity.

In summary, our reported M_5_ mAChR crystal structure has allowed for the comparison of all five mAChR subtypes and has revealed that subtle differences in the ECL regions are a major determinant in ligand selectivity, regardless of the ligand being orthosteric or allosteric. As the M_1_, M_4_, and M_5_ mAChRs continue to emerge as exciting drug targets for the treatment of CNS disorders, it will be important to understand both the structural and dynamic differences between all five mAChR subtypes in order to aid design of safer and more effective small-molecule therapeutics.

## METHODS

### Cloning

The human M_5_ muscarinic receptor gene (cDNA.org) was cloned into a pFastBac vector containing an N-terminal Flag epitope and a C-terminal 10x histidine tag (cDNA.org). TEV cleavage site was introduced in order to remove the Flag epitope and the first 20 amino acids of the protein. Residues 225-430 of ICL3 were removed and replaced with T4L (Supplementary Figure 1). In addition, to stabilize the inactive state we introduced the mutation S117^3.39^R (23, 25). For pharmacology experiments M_2_ and M_5_ mAChR DNA was cloned into a pEF5/FTR/V5 vector (Invitrogen) using the Flp-In-CHO cell system (Invitrogen). To generate the M_2_/M_5_ ECL chimeras overlap extension PCR was used with primers specific to each ECL region. All DNA constructs were sequenced to confirm the correct nucleotide sequence using the Australian Genome Research Facility (Melbourne, Australia).

### Synthesis of the bis-ammonium alkane ligands

4B-C_7_/3-phth and 4P-C_7_/3-phth were synthesized as previously described (27). Synthesis for 4P-C_7_/3-bromo-phth and 4P-aryl C_7_/3-bromo-phth is described in Supplementary Data.

### M5 receptor expression and purification

M_5_-T4L was expressed in Sf9 cells using the Bac-to-Bac Baculovirus Expression System (Invitrogen). Cells were infected at a cell density of 4.0 × 10^6^ cells/millilitre with 10 μM atropine. Cells were harvested 60-72 hours later. All purification steps were performed in the presence of 1 μM triotropium. Insect cells were lysed in a buffer containing 10 mM Tris pH 7.5, 1 mM EDTA, 1 mM MgCl_2_, 1 mg/ml iodoacetamide, benzonase, and protease inhibitors. Cell membranes were then solubilised in 30 mM HEPES pH 7.4, 750 mM NaCl, 30 % glycerol, 1 % dodecyl maltoside (DDM), 0.2 % cholate, 0.03 % cholesterol hemisuccinate (CHS), 1 mM MgCl_2_, 1 mg/ml iodoacetamide, benzonase and protease inhibitors for 90 minutes at 4°C. After removing the insoluble debris, 25 mL of Ni-NTA resin, 1 mg/ml iodoacetamide and protease inhibitors were added and incubated with the protein for 2 hours 4°C. The Ni-NTA resin was pelleted using a table top centrifuge and then washed using 30 mM HEPES pH 7.4, 750 mM NaCl, 30 % glycerol, 5 mM imidazole pH 8.0, 0.1 % DDM, 0.02 % cholate and 0.003 % CHS. The protein was eluted using the same buffer supplemented with 250 mM imidazole. The elution was supplemented with 2 mM CaCl_2_ and then loaded onto an anti-Flag M1 antibody column. The buffer/DDM was exchanged into 30 mM HEPES pH 7.4, 100 mM NaCl, 2 mM CaCl_2_, 0.1 % lauryl maltose neopentyl glycol (LMNG) and 0.01 % CHS over the course of 30 min. The resin was then washed with 10x CMC buffer (30 mM HEPES pH 7.4, 100 mM NaCl, 0.01 % LMNG and 0.001 % CHS) with 2 mM CaCl_2_ before the protein was eluted using the same buffer supplemented with 10 mM EDTA and 0.2 mg/ml FLAG peptide. TEV protease was added (1mg) overnight at 4° before concentrating the protein for size-exclusion chromatography using a Superdex S200 increase column in 10x CMC buffer. Monodispersed fractions were pooled together (Supplementary Figure 2), concentrated to 80 absorbance units (~50 mg/ml) and flash frozen in small aliquots using liquid nitrogen.

### Crystallization and structure determination

Purified M_5_-T4L(S3.39R) bound to tiotropium was crystallized using LCP. For allosteric modulator co-crystallization, the modulator was added to purified protein at a final concentration of 2.5 mM. The sample was incubated on ice for 3 hours before it was mixed into 10:1 (w/w) monoolein:cholesterol in 1:1.5 w/w protein:lipid ratio. LCP crystallization was performed by spotting 25-30 nL of samples on siliconized 96-well glass plate overlaying the samples with 600 nL of precipitant solution using the Gryphon LCP (Art Robbins Instruments). Sealed glass plates were incubated at 20 °C. Crystals appeared in the first 24 hours and grew to full size in the following 1-2 days. The best diffracting crystals grew in 100 mM DL-Malic acid pH 6.0, 220-280 mM ammonium tartrate dibasic and 37-41% PEG 400. For the data collection, whole drops were harvested using mesh grid loops (Mitegen) and flash frozen in liquid nitrogen.

X-ray diffraction data were collected at the SPring-8 (Japan) beamline BL32XU (49) and the MX2 beamline at the Australian Synchrotron (50). Diffraction data at SPring-8 was collected using the automatic data-collection system ZOO (51). Diffraction data was processed using KAMO (52) with XDS (53). The structure was solved using molecular replacement with M_3_-mT4L (4U15) as a search model for the receptor and an ensemble of T4L molecules for T4L. Structure refinement was performed with Phenix (54), and the models were validated with MolProbity (55). Structure figures were prepared with PyMol.

### Pharmacology of crystallization constructs

Sf9 cells expressing M_5_-T4L (S117R) or WT M_5_ mAChR were harvested after 60 hours. Sf9 cell membranes were prepared by homogenization and centrifugation. The final membrane pellet was resuspended in 20 mM HEPES pH 7.4 and 0.1 mM EDTA. Protein concentration was determined by absorbance at 280 nm and membranes were stored at −80 °C. Assays were conducted in UniFilter-96 GF/B plates (PerkinElmer) with 1 ug of membranes per well in a final volume of 300 μl binding buffer consisting of 20 mM HEPES, 100 mM NaCl, and 10 mM MgCl_2_ at pH 7.4. Non-specific binding was defined in the presence of 1 μM atropine. Assays were stopped by vacuum filtration and washed three times with ice-cold 0.9% sodium chloride. Plates were allowed to dry before 40 μL of Microscint-0 (PerkinElmer) was added to each well. Radioactivity was measured on a MicroBeta2 microplate counter. [^3^H]NMS affinity (KA) was determined in saturation binding experiments using 7 different concentrations of [^3^H]NMS (0.03–30 nM). Equilibration was 1 hour at room temperature. Competition binding assays and allosteric modulator assays were performed by incubating membranes with a K_A_ concentration of [^3^H]NMS and varying concentrations of compounds. The reactions were left at room temperature and harvested after 4 hours. Similar competition experiments with tiotropium were performed in the absence and presence of 30 μM of the allosteric modulators.

### Mammalian cell culture

For stable expression, DNA constructs in pEF5/FTR/V5 (ThermoFisher) were transfected into the Flp-In-CHO cell line (ThermoFisher) as previously described (22). A suspected error with the M_2_–M_5_-ECL1 and M_5_–M_2_-all-ECL cell lines led us to resequencing all of the constructs and performing transient transfections for these constructs and also retesting all of the M_2_ cell lines. Cells were maintained in DMEM containing 10% FBS, 16 mM HEPES pH 7.4, and 400 μg ml^−1^ hygromycin B. Mycoplasma testing was performed regularly on cell lines using the MycoAlertTM kit (Lonza); cell lines were mycoplasma-free before experiments were conducted.

### Pharmacology of the M_2_/M_5_ mAChR chimeras

Flp-In-CHO cells either stably or transiently expressing the M_2_/M_5_ mAChR chimeras were seeded in 96-well Isoplates (PerkinElmer Life Sciences) at a concentration of 20,000 – 25,000 cells per well a day before the experiment was performed in a humidified atmosphere at 37°C, 5% CO_2_. The next day, cells were washed with assay buffer consisting of 110 mM NaCl, 5.4 mM KCl, 1.8 mM CaCl_2_, 1 mM MgSO_4_, 25 mM glucose, 50 mM HEPES, and 58 mM sucrose, pH 7.4. Saturation binding experiments were performed similar to above with assay volumes of 150 μL. Kinetic binding dissociation experiments were performed by incubating cells with 0.2 nM [^3^H]NMS for 3 hours at room temperature before a reverse time-course was performed. Dissociation was initiated by atropine and treatments consisted of 10 μM atropine and either vehicle, 10 μM ML375, or 10 μM 4P-C_7_/3-phth being added at the indicated time points. At the end of the time course, radioligand was removed by inverting the plate and followed by 3 washes with ice-cold 0.9% sodium chloride. Bound radioactivity was assessed by liquid scintillation using Optiphase Supermix (100 μL) and counting on a MicroBeta2 Plate Counter.

### Data Analysis

Data were analysed using Prism 8.2 (GraphPad). Saturation binding curves were fitted to a one site binding curve accounting for total and non-specific binding. Competition binding curves were fitted to a one-site binding inhibition model. Allosteric titrations between [^3^H]NMS and modulators were fit to an allosteric ternary complex model. Radioligand dissociation data were fitted to a mono-exponential decay function.

## Author Contributions

Z.V., P.R.G., and D.M.T. performed cloning, protein expression, purification, crystallization, data collection, structure refinement, and radioligand binding experiments on the crystallization constructs. K.H. collected and processed data all the data from SPring-8. S.J. and J.B. designed and performed the synthesis of 4P-C_7_/3-bromo-phth and 4P-aryl-C_7_/3-bromo-phth. R.R. and J.B. designed and performed the synthesis of 4P-C_7_/3-phth and 4B-C_7_/3-phth. For the M_2_/M_5_ mAChR chimera experiments: Z.V. performed cloning and generated the stable cell lines. Molecular pharmacology experiments were performed by A.E.B., Z.V., E.T.W., G.T., and W.A.C.B. C.L. provided ML375. C.V., C.L., J.B., A.B.T., P.M.S., A.C., and D.M.T. provided overall project design and supervision. Z.V., P.R.G., A.C., and D.M.T. wrote the manuscript with contributions from all authors.

## Acknowledgements

The synchrotron radiation experiments were performed at the BL32XU beamline at SPring-8 with the approval of JASRI (Proposal No. 2017B2731) and the MX2 beamline at the Australian Synchrotron (CAP13670). This work was funded by a Wellcome Trust Collaborative Award (201529/Z/16/Z), and supported by National Health and Medical Research Council of Australia (NHMRC) project (APP1138448) and program (APP1055134) grants. J.B. is a NHMRC Principal Research Fellow, P.M.S. and A.C. are NHMRC Senior Principal Research Fellows, and D.M.T. is an Australian Research Council Discovery Early Career Research Fellow.

## Uncategorized References

1. M. P. Caulfield, N. J. Birdsall, International Union of Pharmacology. XVII. Classification of muscarinic acetylcholine receptors. Pharmacol Rev 50, 279–290 (1998).

2. A. C. Kruse, J. Hu, B. K. Kobilka, J. Wess, Muscarinic acetylcholine receptor X-ray structures: potential implications for drug development. Curr Opin Pharmacol 16, 24–30 (2014).

3. R. P. Yasuda et al., Development of antisera selective for m4 and m5 muscarinic cholinergic receptors: distribution of m4 and m5 receptors in rat brain. Mol Pharmacol 43, 149–157 (1993).

4. A. M. Bender, A. T. Garrison, C. W. Lindsley, The Muscarinic Acetylcholine Receptor M5: Therapeutic Implications and Allosteric Modulation. ACS Chem Neurosci 10, 1025–1034 (2019).

5. G. L. Forster, C. D. Blaha, Laterodorsal tegmental stimulation elicits dopamine efflux in the rat nucleus accumbens by activation of acetylcholine and glutamate receptors in the ventral tegmental area. Eur J Neurosci 12, 3596–3604 (2000).

6. G. L. Forster, C. D. Blaha, Pedunculopontine tegmental stimulation evokes striatal dopamine efflux by activation of acetylcholine and glutamate receptors in the midbrain and pons of the rat. Eur J Neurosci 17, 751–762 (2003).

7. G. L. Forster, J. S. Yeomans, J. Takeuchi, C. D. Blaha, M5 muscarinic receptors are required for prolonged accumbal dopamine release after electrical stimulation of the pons in mice. J Neurosci 22, RC190 (2002).

8. S. Steidl, A. D. Miller, C. D. Blaha, J. S. Yeomans, M(5) muscarinic receptors mediate striatal dopamine activation by ventral tegmental morphine and pedunculopontine stimulation in mice. PLoS One 6, e27538 (2011).

9. A. Elhusseiny, Z. Cohen, A. Olivier, D. B. Stanimirovic, E. Hamel, Functional acetylcholine muscarinic receptor subtypes in human brain microcirculation: identification and cellular localization. J Cereb Blood Flow Metab 19, 794–802 (1999).

10. S. K. Tayebati, M. A. Di Tullio, D. Tomassoni, F. Amenta, Localization of the m5 muscarinic cholinergic receptor in rat circle of Willis and pial arteries. Neuroscience 122, 205–211 (2003).

11. R. Araya et al., Loss of M5 muscarinic acetylcholine receptors leads to cerebrovascular and neuronal abnormalities and cognitive deficits in mice. Neurobiol Dis 24, 334–344 (2006).

12. M. Yamada et al., Cholinergic dilation of cerebral blood vessels is abolished in M(5) muscarinic acetylcholine receptor knockout mice. Proc Natl Acad Sci U S A 98, 14096–14101 (2001).

13. A. S. Basile et al., Deletion of the M5 muscarinic acetylcholine receptor attenuates morphine reinforcement and withdrawal but not morphine analgesia. Proc Natl Acad Sci U S A 99, 11452–11457 (2002).

14. A. Fink-Jensen et al., Role for M5 muscarinic acetylcholine receptors in cocaine addiction. J Neurosci Res 74, 91–96 (2003).

15. M. Thomsen et al., Reduced cocaine self-administration in muscarinic M5 acetylcholine receptor-deficient mice. J Neurosci 25, 8141–8149 (2005).

16. A. E. Berizzi et al., Muscarinic M5 receptors modulate ethanol seeking in rats. Neuropsychopharmacology 43, 1510–1517 (2018).

17. R. W. Gould et al., Acute Negative Allosteric Modulation of M5 Muscarinic Acetylcholine Receptors Inhibits Oxycodone Self-Administration and Cue-Induced Reactivity with No Effect on Antinociception. ACS Chem Neurosci 10.1021/acschemneuro.9b00274 (2019).

18. B. W. Gunter et al., Selective inhibition of M5 muscarinic acetylcholine receptors attenuates cocaine self-administration in rats. Addict Biol 23, 1106–1116 (2018).

19. P. R. Gentry et al., Discovery of the first M5-selective and CNS penetrant negative allosteric modulator (NAM) of a muscarinic acetylcholine receptor: (S)-9b-(4-chlorophenyl)-1-(3,4-difluorobenzoyl)-2,3-dihydro-1H-imidazo[2,1-a]isoi ndol-5(9bH)-one (ML375). J Med Chem 56, 9351–9355 (2013).

20. D. A. Sykes et al., The Influence of receptor kinetics on the onset and duration of action and the therapeutic index of NVA237 and tiotropium. J Pharmacol Exp Ther 343, 520–528 (2012).

21. A. C. Kruse et al., Structure and dynamics of the M3 muscarinic acetylcholine receptor. Nature 482, 552–556 (2012).

22. D. M. Thal et al., Crystal structures of the M1 and M4 muscarinic acetylcholine receptors. Nature 531, 335–340 (2016).

23. Y. Kajiwara, S. Yasuda, Y. Takamuku, T. Murata, M. Kinoshita, Identification of thermostabilizing mutations for a membrane protein whose three-dimensional structure is unknown. J Comput Chem 38, 211–223 (2017).

24. J. A. Ballesteros, H. Weinstein, “[19] Integrated methods for the construction of three-dimensional models and computational probing of structure-function relations in G protein-coupled receptors” in Methods in Neurosciences, S. C. Sealfon, Ed. (Academic Press, 1995), vol. 25, pp. 366–428.

25. R. Suno et al., Structural insights into the subtype-selective antagonist binding to the M2 muscarinic receptor. Nat Chem Biol 14, 1150–1158 (2018).

26. X. P. Huang, S. Prilla, K. Mohr, J. Ellis, Critical amino acid residues of the common allosteric site on the M2 muscarinic acetylcholine receptor: more similarities than differences between the structurally divergent agents gallamine and bis(ammonio)alkane-type hexamethylene-bis-[dimethyl-(3-phthalimidopropyl)ammonium]dibromide. Mol Pharmacol 68, 769–778 (2005).

27. R. O. Dror et al., Structural basis for modulation of a G-protein-coupled receptor by allosteric drugs. Nature 503, 295–299 (2013).

28. T. S. Thorsen, R. Matt, W. I. Weis, B. K. Kobilka, Modified T4 Lysozyme Fusion Proteins Facilitate G Protein-Coupled Receptor Crystallogenesis. Structure 22, 1657–1664 (2014).

29. W. Bender, M. Staudt, C. Trankle, K. Mohr, U. Holzgrabe, Probing the size of a hydrophobic binding pocket within the allosteric site of muscarinic acetylcholine M2-receptors. Life Sci 66, 1675–1682 (2000).

30. K. Haga et al., Structure of the human M2 muscarinic acetylcholine receptor bound to an antagonist. Nature 482, 547–551 (2012).

31. E. Jurrus et al., Improvements to the APBS biomolecular solvation software suite. Protein Sci 27, 112–128 (2018).

32. D. M. Thal, A. Glukhova, P. M. Sexton, A. Christopoulos, Structural insights into G-protein-coupled receptor allostery. Nature 559, 45–53 (2018).

33. A. Bock, R. Schrage, K. Mohr, Allosteric modulators targeting CNS muscarinic receptors. Neuropharmacology 136, 427–437 (2018).

34. K. J. Gregory, P. M. Sexton, A. Christopoulos, Allosteric modulation of muscarinic acetylcholine receptors. Curr Neuropharmacol 5, 157–167 (2007).

35. W. A. C. Burger, P. M. Sexton, A. Christopoulos, D. M. Thal, Toward an understanding of the structural basis of allostery in muscarinic acetylcholine receptors. J Gen Physiol 150, 1360–1372 (2018).

36. S. Buller, D. P. Zlotos, K. Mohr, J. Ellis, Allosteric site on muscarinic acetylcholine receptors: a single amino acid in transmembrane region 7 is critical to the subtype selectivities of caracurine V derivatives and alkane-bisammonium ligands. Mol Pharmacol 61, 160–168 (2002).

37. A. L. Gnagey, M. Seidenberg, J. Ellis, Site-directed mutagenesis reveals two epitopes involved in the subtype selectivity of the allosteric interactions of gallamine at muscarinic acetylcholine receptors. Mol Pharmacol 56, 1245–1253 (1999).

38. S. Prilla, J. Schrobang, J. Ellis, H. D. Holtje, K. Mohr, Allosteric interactions with muscarinic acetylcholine receptors: complex role of the conserved tryptophan M2422Trp in a critical cluster of amino acids for baseline affinity, subtype selectivity, and cooperativity. Mol Pharmacol 70, 181–193 (2006).

39. U. Voigtlander et al., Allosteric site on muscarinic acetylcholine receptors: identification of two amino acids in the muscarinic M2 receptor that account entirely for the M2/M5 subtype selectivities of some structurally diverse allosteric ligands in N-methylscopolamine-occupied receptors. Mol Pharmacol 64, 21–31 (2003).

40. J. R. Lane, L. T. May, R. G. Parton, P. M. Sexton, A. Christopoulos, A kinetic view of GPCR allostery and biased agonism. Nat Chem Biol 13, 929–937 (2017).

41. A. C. Kruse et al., Muscarinic acetylcholine receptors: novel opportunities for drug development. Nat Rev Drug Discov 13, 549–560 (2014).

42. A. K. Apel et al., Crystal Structure of CC Chemokine Receptor 2A in Complex with an Orthosteric Antagonist Provides Insights for the Design of Selective Antagonists. Structure 27, 427–438.e425 (2019).

43. H. Liu et al., Structure-guided development of selective M3 muscarinic acetylcholine receptor antagonists. Proc Natl Acad Sci U S A 115, 12046–12050 (2018).

44. J. D. McCorvy et al., Structure-inspired design of beta-arrestin-biased ligands for aminergic GPCRs. Nat Chem Biol 14, 126–134 (2018).

45. A. Strasser, H. J. Wittmann, R. Seifert, Binding Kinetics and Pathways of Ligands to GPCRs. Trends Pharmacol Sci 38, 717–732 (2017).

46. D. C. Swinney, B. A. Haubrich, I. Van Liefde, G. Vauquelin, The Role of Binding Kinetics in GPCR Drug Discovery. Curr Top Med Chem 15, 2504–2522 (2015).

47. R. A. Copeland, The drug-target residence time model: a 10-year retrospective. Nat Rev Drug Discov 15, 87–95 (2016).

48. C. S. Tautermann et al., Molecular basis for the long duration of action and kinetic selectivity of tiotropium for the muscarinic M3 receptor. J Med Chem 56, 8746–8756 (2013).

49. K. Hirata et al., Achievement of protein micro-crystallography at SPring-8 beamline BL32XU. Journal of Physics: Conference Series 425, 012002 (2013).

50. D. Aragao et al., MX2: a high-flux undulator microfocus beamline serving both the chemical and macromolecular crystallography communities at the Australian Synchrotron. J Synchrotron Radiat 25, 885–891 (2018).

51. K. Hirata et al., ZOO: an automatic data-collection system for high-throughput structure analysis in protein microcrystallography. Acta Crystallogr D Struct Biol 75, 138–150 (2019).

52. K. Yamashita, K. Hirata, M. Yamamoto, KAMO: towards automated data processing for microcrystals. Acta Crystallogr D Struct Biol 74, 441–449 (2018).

53. W. Kabsch, Xds. Acta Crystallogr D Biol Crystallogr 66, 125–132 (2010).

54. P. D. Adams et al., PHENIX: a comprehensive Python-based system for macromolecular structure solution. Acta Crystallogr D Biol Crystallogr 66, 213–221 (2010).

55. V. B. Chen et al., MolProbity: all-atom structure validation for macromolecular crystallography. Acta Crystallogr D Biol Crystallogr 66, 12–21 (2010).

