## Supplementary Data for "Crystal structure of the M_5_ muscarinic acetylcholine receptor"

### Supplementary Data for
### Crystal structure of the $M_5$ muscarinic acetylcholine receptor

Ziva Vuckovic[1]*, Patrick R. Gentry[1]*, Alice E. Berizzi[1], Kunio Hirata[2], Swapna Varghese[3], Geoff Thompson[1], Emma T. van der Westhuizen[1], Wessel A.C. Burger[1], Raphaël Rahmani[3], Celine Valant[1], Christopher J. Langmead[1], Craig W. Lindsley[4], Jonathan Baell[3], Andrew B. Tobin[5], Patrick M. Sexton[1,6], Arthur Christopoulos[1¶] and David M. Thal[1¶]

1. Drug Discovery Biology and Department of Pharmacology, Monash Institute of Pharmaceutical Sciences, Monash University, Parkville 3052, Victoria, Australia

2. RIKEN, SPring-8 Center, Hyogo, Japan

3. Medicinal Chemistry, Monash Institute of Pharmaceutical Sciences, Monash University, Parkville 3052, Victoria, Australia

4. Departments of Pharmacology & Chemistry, Vanderbilt Center for Neuroscience Drug Discovery, Vanderbilt University, Nashville, Tennessee 37232, United States.

5. Centre for Translational Pharmacology, Institute of Molecular, Cell and Systems Biology, College of Medical, Veterinary and Life Sciences, University of Glasgow, Glasgow G12 8QQ, United Kingdom.

6. School of Pharmacy, Fudan University, Shanghai 201203, China.

*These authors contributed equally to this work.
¶ To whom correspondence should be addressed:

David Thal,

Arthur Christopoulos,

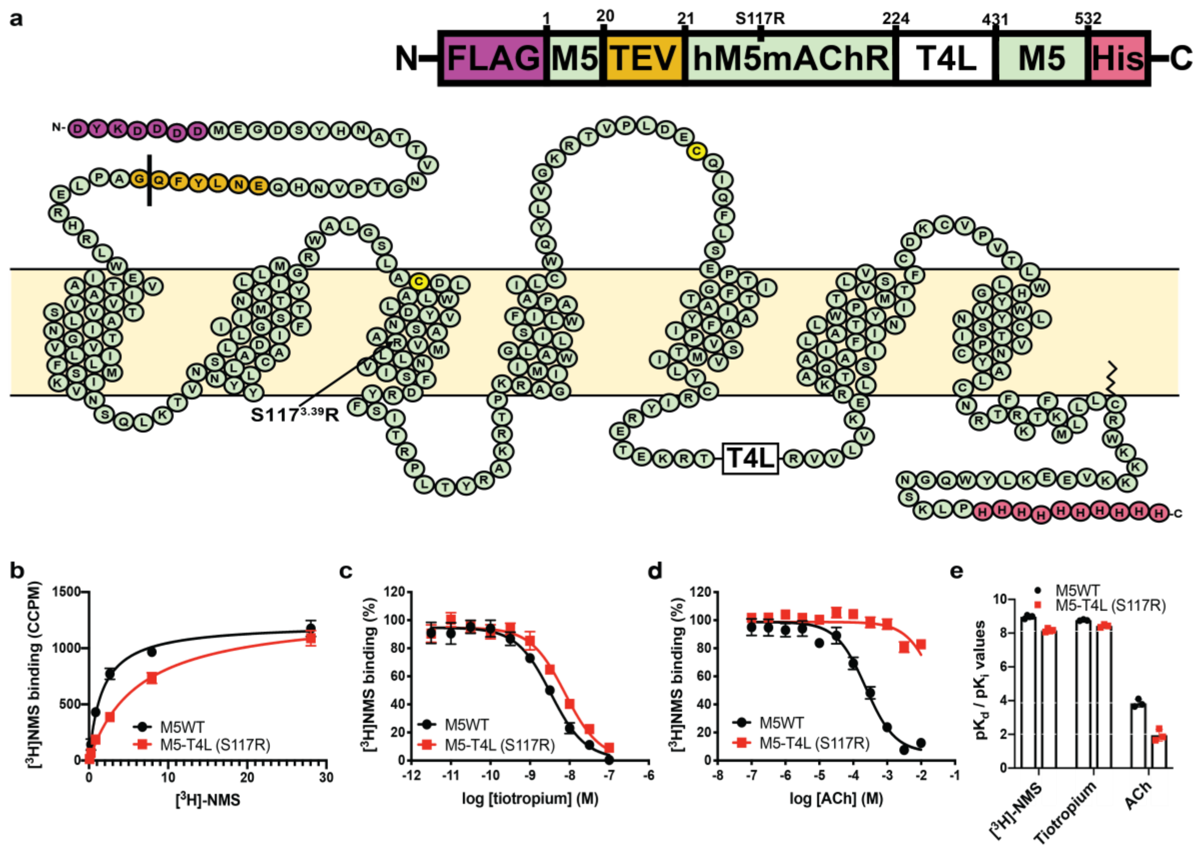


**Supplementary Figure 1. Characterization of the M$_5$-T4L (S117R) construct. (a)** Crystallization construct used for the M$_5$ mAChR structure. **(b)** Saturation [$^3$H]-NMS and **(c,e)** [$^3$H]-NMS equilibrium competition binding of **(c)** tiotropium or **(d)** acetylcholine (ACh) on WT M$_5$ or M$_5$-T4L(S117R) from Sf9 cell membranes. **(e)** Mean ± S.E.M. values with individual pK$_d$ or pK$_i$ values are shown.

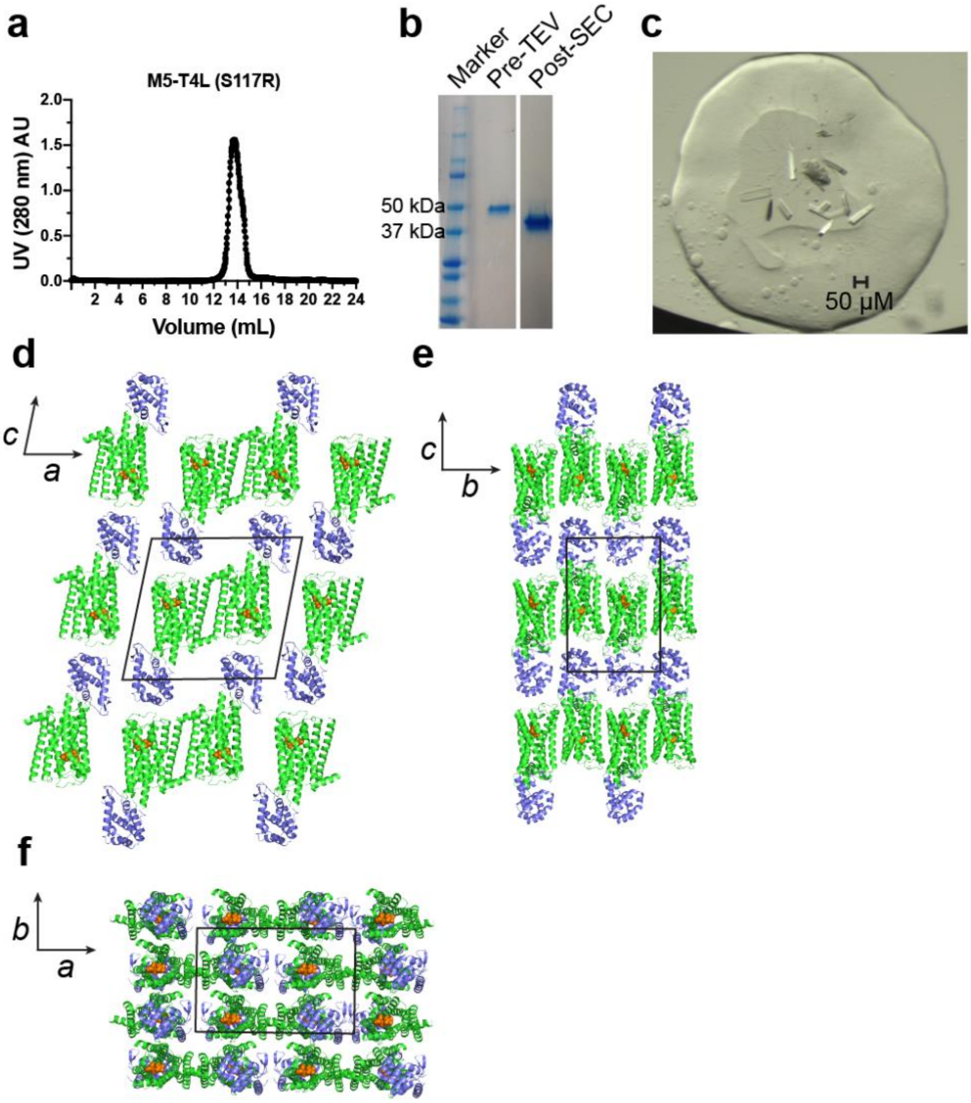


**Supplementary Figure 2. Purification and crystallization of M5-T4L (S117R). (a)** A size exclusion chromatography chromatogram of the purified TEV treated receptor. (**b**) Coomassie stained SDS-PAGE of M5-T4L(S117R) before TEV cleavage and after SEC purification. (**c**) M5-T4L(S117R)•tiotropium with 1 mM 4B-$C_7$/3-phth was setup into LCP. Crystals of approximately 100 µm grew in conditions consisting of: 100 mM DL-Malic acid pH 6.0, 220-280 mM ammonium tartrate dibasic and 37-41% PEG 400. (d-f) Crystal packing of the M5-T4L(S117R) with views along the **(d)** b-axis, **(e)** a-axis, and **(f)** c-axis with the receptor colored in green and T4L in blue.

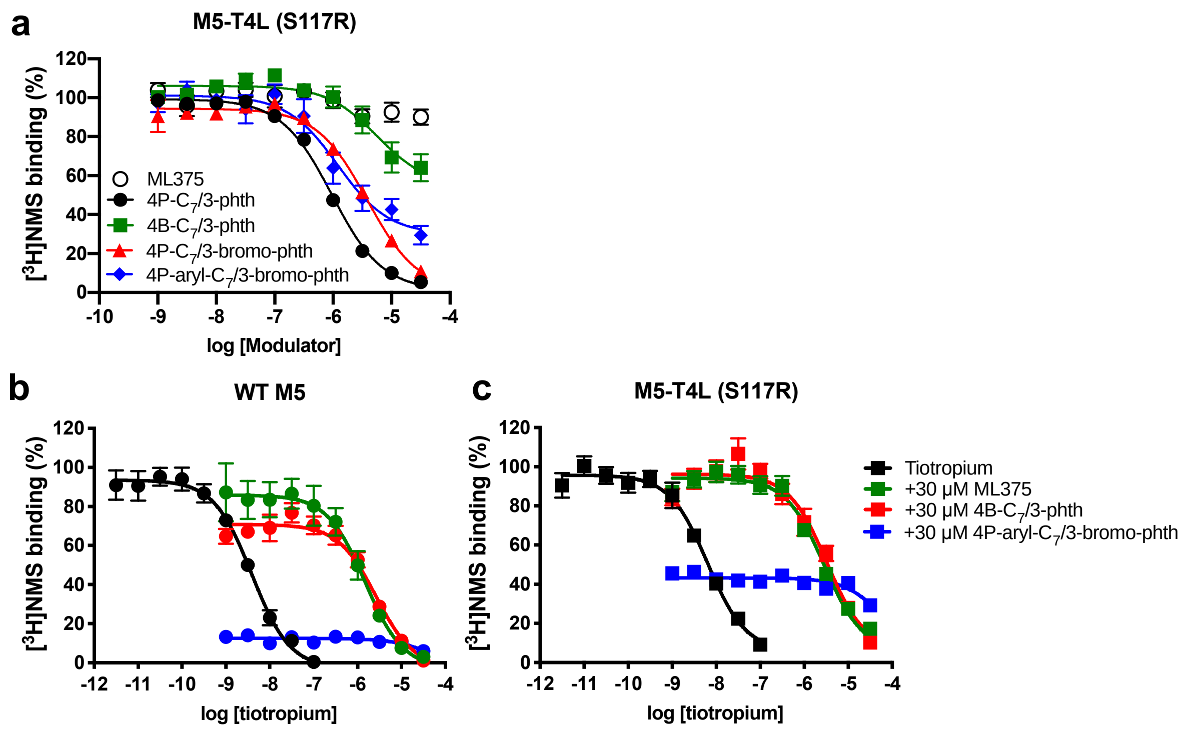


**Supplementary Figure 3. Characterization of the NAMs used in this study again M5-T4L (S117R). (a)** Equilibrium competition binding between [$^{3}$H]NMS and the indicated NAMS. ML375 did not show appreciable competition and was not analysed. Quantitative values are listed in Supplementary Table 2. Equilibrium competition between [$^{3}$H]NMS and tiotropium in the presence of 30 µM of the allosteric modulators used in the X-ray crystallography experiments tested against (**b**) wild type (WT) M5 and (**c**) M5-T4L (S117R). Data shown are the mean ± S.E.M. of three or more independent experiments performed in duplicate.

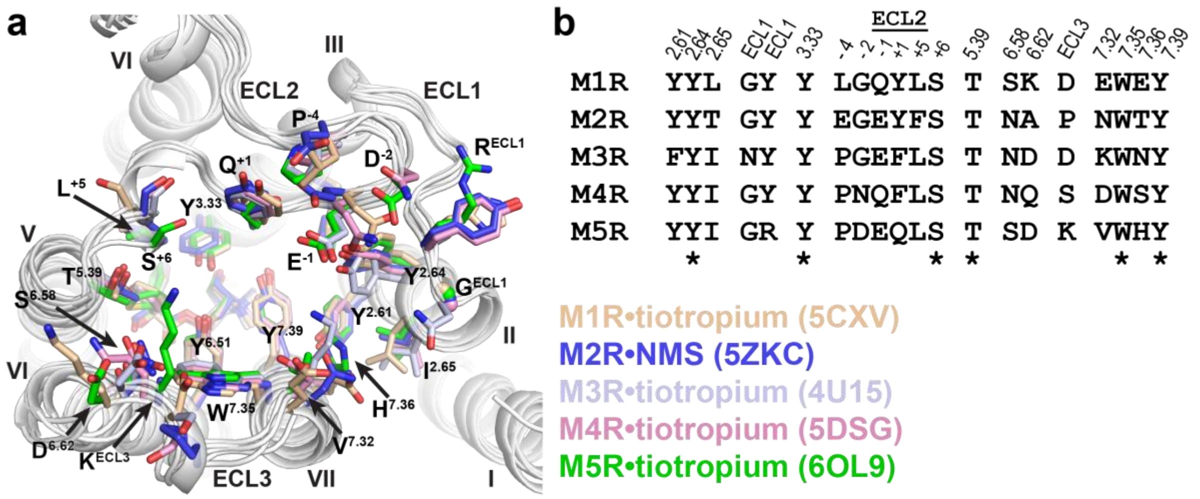


**Supplementary Figure 4. Comparison of the ECV across all five mAChR subtypes.** (**a**) Structures from Figure 2b are shown with the receptor cartoons in white, and individual residues in the ECV coloured according to subtype. Amino acids are labelled according to the $M_5$ mAChR subtype. Residues in ECL2 are numbered relative to the position of the cysteine residue in the conserved disulfide bond between ECL2 and TM3. (**b**) Comparison of the amino acid identities for ECV residues across all five mAChR subtypes. Conserved residues are indicated with an asterix.

**Supplementary Table 1 | Data collection and refinement statistics**

| **Data collection** | M5-T4L•tiotropium | M5-T4L•tiotropium[a] (PDB: 6OL9) | M5-T4L•tiotropium[b] | M5-T4L•tiotropium[c] | M5-T4L•tiotropium[d] |
|---|---|---|---|---|---|
| Beamline | Spring-8, RIKEN BL32XU | Spring-8, RIKEN BL32XU | Spring-8, RIKEN BL32XU | Spring-8, RIKEN BL32XU | Spring-8, RIKEN BL32XU |
| Number of crystals | 130 | 318 | 316 | 300 | 2 |
| Space group | C 1 2 1 | C 1 2 1 | C 1 2 1 | C 1 2 1 | C 1 2 1 |
| Cell dimensions | | | | | |
| *a, b, c* (Å) | 97.5, 63.2, 91.2 | 96.8, 63.6, 91.9 | 96.9, 63.3, 91.7 | 96.3, 63.3, 91.3 | 96.46, 63.48, 91.62 |
| α, β, γ (°) | 90, 101.7, 90 | 90, 102.4, 90 | 90, 102.2, 90 | 90, 102.2, 90 | 90, 103.0, 90 |
| Resolution (Å)[e] | 47.80 – 3.10 (3.31 – 3.10) | 48.07 – 2.55 (2.66 – 2.55) | 47.90 – 2.50 (2.60 – 2.50) | 47.77 – 2.70 (2.83– 2.70) | 48.04 – 2.59 (2.71 – 2.59) |
| $R_{merge}$ (%) | 41.6 (>100) | 30.2 (>100) | 44.8 (>100) | 40.2 (>100) | 9.4 (24.1) |
| Rpim (%) | 11.9 (107.4) | 3.9 (53.3) | 5.6 (29.7) | 5.4 (42.8) | 6.7 (20.6) |
| <I/σI> | 4.9 (0.6) | 13.6 (1.8) | 11.1 (0.9) | 10.8 (2.0) | 8.4 (2.1) |
| $CC_{1/2}$ (%) | 98.3 (33.8) | 99.2 (68.4) | 99.7 (13.5) | 99.6 (71.2) | 95.5 (88.3) |
| Completeness | 98.6 (98.5) | 100 (100) | 100 (100) | 100 (100) | 75.2 (40.2) |
| Multiplicity | 10.1 (10.1) | 56.5 (41.9) | 52.3 (12.4) | 51.2 (40.1) | 2.4 (1.7) |
| **Refinement** | | | | | |
| Resolution (Å) | 47.80 - 3.40 | 48.07 - 2.55 | 47.90 - 2.50 | 47.77-2.70 | 48.04 – 2.59 |
| No. of reflections working / test set | 13,552 / 680 | 17,926 / 882 | 18,391 / 906 | 14,904 / 725 | 20,921 / 1009 |
| $R_{work}/R_{free}$ (%) | 26.7 / 29.7 | 23.0/ 25.4 | 24.0 / 26.1 | 23.1 / 25.0 | 22.6 / 25.3 |
| No. of atoms | | | | | |
| Protein | 3127 | 3281 | 3275 | 3278 | 3275 |
| Ligands | 70 | 105 | 105 | 126 | 105 |
| Average B-factors (Å$^2$) | 103.0 | 96.0 | 89.6 | 92.9 | 80.5 |
| Receptor | 88.9 | 81.6 | 74.1 | 77.9 | 70.5 |
| T4 lysozyme | 145.1 | 127.5 | 124.2 | 126.5 | 102.5 |
| Tiotropium | 58.9 | 57.7 | 50.5 | 51.7 | 52.9 |
| Allosteric site PEG 400 | 75.2 | 107.2 | 85.6 | 90.3 | 88.0 |
| Waters | 29.9 | 65.0 | 53.9 | 63.7 | 61.3 |
| Other ligands | 69.0 | 94.4 | 83.5 | 87.1 | 84.8 |
| RMS deviation from ideality | | | | | |
| Bond length (Å) | 0.002 | 0.002 | 0.002 | 0.003 | 0.004 |
| Bond angles (°) | 0.55 | 0.51 | 0.45 | 0.69 | 0.78 |
| Ramachandran statistics[f] | | | | | |
| Favored regions (%) | 98.7 | 98.5 | 98.8 | 98.8 | 98.05 |
| Allowed regions (%) | 1.3 | 1.5 | 1.2 | 1.2 | 1.70 |
| Outliers (%) | 0.0 | 0.0 | 0.0 | 0.0 | 0.24 |

[a-d] Co-crystallized with [a]4B-$C_7$/3-phth, [b]4P-aryl-$C_7$/3-bromo-phth, [c]ML375, [d]4P-$C_7$/3-bromo-phth.
[e] Highest shell statistics in parenthesis.
[f] As calculated by Molprobity.

**Supplementary Table 2 | Binding affinity values for allosteric modulators against M5-T4L(S117R) from Sf9 membranes.**

| Ligand | pKb |
|---|---|
| 4P-$C_7$/3-phth | 6.3 ± 0.01 |
| 4B-$C_7$/3-phth | 5.4 ± 0.1 |
| 4P-$C_7$/3-bromo-phth | 5.7 ± 0.07 |
| 4P-aryl-$C_7$/3-bromo-phth | 6.1 ± 0.2 |
| ML375 | N.D. |

Data represent the mean ± S.E.M. of three or more independent experiments performed in duplicate. N.D. Value not determined from curve fit.

**Supplementary Table 3 | [$^3$H]NMS Equilibrium binding parameter estimates.**

| Constructs | pKd | Bmax (fmol/10^5 cells) |
|---|---|---|
| M2 WT | 9.6 ± 0.2 | 12.8 ± 2.9 |
| M2–M5-ECL1 | 9.3 ± 0.1 | 4.6 ± 1.2 # |
| M2–M5-ECL2 | 9.3 ± 0.1 | 13.0 ± 0.6 |
| M2–M5-ECL3 | 10.0 ± 0.03 | 14.3 ± 1.8 |
| M2–M5-all-ECLs | 9.4 ± 0.04 | 18.0 ± 2.0 |
| M5 WT | 9.4 ± 0.2 | 13.8 ± 1.2 |
| M5–M2-ECL1 | 9.3 ± 0.02 | 17.0 ± 0.6 |
| M5–M2-ECL2 | 9.7 ± 0.02 | 7.3 ± 0.7 * |
| M5–M2-ECL3 | 9.7 ± 0.1 | 8.4 ± 1.3 * |
| M5–M2-all-ECLs | 9.0 ± 0.1 | 3.0 ± 0.4 # |

Data represent the mean ± S.E.M. of three or more independent experiments performed in duplicate. *significantly different from WT, $p < 0.05$, one-way ANOVA, Dunnett's post hoc test. #Constructs were tested using transient transfections, hence the comparatively lower expression values.

**Supplementary Table 4 | Effects of M2 and M5 chimeras on the kinetics of [$^3$H]NMS dissociation.**

| | [$^3$H]NMS (control) | | + 10 µM ML375 | | + 10 µM 4P-C7/3-phth | |
|---|---|---|---|---|---|---|
| Constructs | $k_{off}$ (min-1) | $t_{1/2}$ (min) | $k_{off}$ (min-1) | $t_{1/2}$ (min) | $k_{off}$ (min-1) | $t_{1/2}$ (min) |
| M2 WT | 0.16 ± 0.04 | 5.7 ± 1.2 | 0.19 ± 0.04 | 4.3 ± 0.8 | < 0.002 [a] | > 300 |
| M2–M5-ECL1 | 0.04 ± 0.002 | 16.4 ± 0.6 | 0.056 ± 0.007 | 12.6 ± 1.4 | < 0.002 [a] | > 300 |
| M2–M5-ECL2 | 0.15 ± 0.007 | 4.6 ± 0.2 | 0.34 ± 0.09 | 2.8 ± 0.6 | 0.02 ± 0.002 | 35.4 ± 3.1 |
| M2–M5-ECL3 | 0.042 ± 0.003 | 16.9 ± 1.4 | 0.057 ± 0.006 | 12.6 ± 1.0 | < 0.002 [a] | > 300 |
| M2–M5-all-ECLs | 0.024 ± 0.0006 | 28.4 ± 0.7 | 0.04 ± 0.006 | 18.7 ± 2.1 | 0.02 ± 0.0006 | 34.9 ± 1.1 |
| M5 WT | 0.0073 ± 0.0008 | 100 ± 11.6 | 0.0043 ± 0.0002 | 162 ± 6.9 | 0.0065 ± 0.0008 | 117 ± 15.9 |
| M5–M2-ECL1 | 0.038 ± 0.007 | 19.7 ± 4.2 | 0.020 ± 0.001 | 34.4 ± 1.7 | 0.013 ± 0.002 | 55.4 ± 10.1 |
| M5–M2-ECL2 | 0.0062 ± 0.0008 | 117 ± 15.4 | 0.0042 ± 0.0003 | 165 ± 9.2 | 0.0022 ± 0.0002 | 319 ± 33.2 |
| M5–M2-ECL3 | 0.017 ± 0.0009 | 40.8 ± 2.0 | 0.008 ± 0.0003 | 90.8 ± 3.3 | 0.012 ± 0.0006 | 57.9 ± 3.1 |
| M5–M2-all-ECLs | 0.036 ± 0.0006 | 19.2 ± 0.3 | 0.034 ± 0.002 | 20.5 ± 1.3 | < 0.002 [a] | > 300 |

Data represent the mean ± S.E.M. of three or more independent experiments performed in duplicate. $t_{1/2}$ is the half-life of dissociation. All experiments were performed in the presence of 10 µM atropine.

[a] A precise value for $k_{off}$ cannot be calculated due to the substantially decreased rate of [$^3$H]NMS dissociation over the time course.

**Synthesis of the bis-ammonium alkane ligands.**

***Materials and Methods for Organic Synthesis.*** Solvents were of analytical grade: ethyl acetate (EtOAc); dichloromethane (DCM); dimethyl formamide (DMF); methanol (MeOH); tetrahydrofuran (THF). Analytical TLC was performed on silica gel 60/$F_{254}$ pre-coated aluminium sheets (0.25 mm, Merck). Flash column chromatography was carried out with silica gel 60, 0.63–0.20 mm (70–230 mesh, Merck); preparative TLC (2 mm, Merck) was performed in 15% MeOH/DCM. Preparative reversed-phase HPLC was performed on an Agilent 1260 Infinity Prep LC (G1361A pump, G2260A autosampler, G1364B fraction collector, and G1315D detector [254 nm]). LC conditions: Alltinea C8 column (5 µm, 250 × 22 mm) using binary gradient elution (solvent A: 0.1% TFA/95.4% $H_2O$/5% $CH_3CN$; solvent B: 0.1% TFA/99.9% $CH_3CN$; 0 to 100% B [9 min], 100% B [1 min]), 30 °C, 20 ml $min^{-1}$); 200-µl injection volume. $^{1}H$, $^{19}F$, and $^{13}C$ nuclear magnetic resonance (NMR) spectra were recorded at 400, 500 MHz or 100 MHz (as specified). Chemical shifts (δ, ppm) are reported relative to the solvent peak ($CDCl_3$: 7.26 [$^{1}H$] or 77.16 [$^{13}C$], DMSO-$d_6$: 2.50 [$^{1}H$] or 39.52 [$^{13}C$]). Proton resonances are annotated as: chemical shift (ppm), multiplicity (br, broad; s, singlet; d, doublet; t, triplet; q, quartet; m, multiplet), coupling constant (*J*, Hz), and number of protons. Analytical HPLC was acquired on an Agilent 1260 Infinity analytical HPLC coupled with a G1322A degasser, G1312B binary pump, G1367E high performance autosampler, G4212B diode array detector. Conditions: Zorbax Eclipse Plus C18 Rapid resolution column (4.6 × 100 mm) with UV detection at 254 nm and 214 nm, 30°C; sample was eluted using a gradient of 5 – 100% of solvent B in solvent A where solvent A: 0.1% aq. TFA, and solvent B: 0.1% TFA in $CH_3CN$ (5 to 100% B [9 min], 100% B [1 min]; 0.5 mL/min). High resolution MS was performed on a Waters Autospec instrument (for compounds **3 - 20**) or an Agilent 6224 TOF LCMS coupled to an Agilent 1290 Infinity LC. All data were acquired and reference mass corrected via a dual-spray electrospray ionization (ESI) source. LC conditions: Agilent Zorbax SB-C18 Rapid Resolution HT (2.1 × 50 mm, 1.8 µm column), 30°C; sample (5 µL) was eluted using a binary gradient (solvent A: 0.1% aq. $HCO_2H$; solvent B: 0.1% $HCO_2H$ in $CH_3CN$; 5 to 100% B [3.5 min], 0.5 mL/min). Purity was determined by HPLC or $^{1}H$ NMR analysis and was found to be ≥95% unless stated otherwise.

**General procedure A:** ***bis-amide formation***

Bis-carboxylic acid was dissolved in 1 M DCM with few drops of DMF and cooled to 0 °C. Oxalyl chloride (2.2 eq.) was then added dropwise and the solution stirred at 0 °C for 2h. The solvent was then evaporated and the residue dissolved in THF and cooled to 0 °C. A solution of dipropyl amine (2.5 eq.) and $Et_3N$ (3 eq.) in THF were then added dropwise and the solution stirred at 0 °C for 30 min and for 2 h at 25 °C. After completion of reaction, 2 M HCl was added to quench the reaction and the organic layer extracted using EtOAc (×3). The combined organic layer was washed with 10% aq. $Na_2CO_3$ and brine, dried with $MgSO_4$ and evaporated to dryness to obtain the desired product.

**General procedure B:** ***bis-amine formation***

To a solution of 1.5 M THF and $LiAlH_4$ (2.6 eq.) at -78 °C, was slowly added bis-amide (1 eq.) dissolved in 1.5 M THF. The reaction was stirred at -78 °C for 2 h and then at 25 °C for 2 h. After completion of reaction, 5 N NaOH was added dropwise till gas evolution ceased. The organic layer was extracted using EtOAc (×3), dried using $MgSO_4$ and evaporated to dryness to obtain the desired product. The compound was further purified by dissolving the crude in ether, followed by acidifying the solution using aq. 1 M HCl. The solution was washed 3 times with ether and the organic layer discarded. The aqueous layer was then basified with aq. 1 N NaOH and extracted with EtOAc (×3). The combined organic layer was then dried using $MgSO_4$ and evaporated to dryness.

**General procedure C:** ***bis-amine quaternarization***

A mixture of bis-amine (1 eq.) and 5-bromo-2-(3-bromopropyl)isoindoline-1,3-dione (2 eq.) in ethylene carbonate (17 eq.) was taken in a sealed tube, flushed with nitrogen gas and heated to 90 °C for 60 h. *Purification method A*: The crude mixture was dissolved in MeOH (1 mL) and purified by preparative TLC eluting with 15% MeOH/DCM. *Purification method B*: The compound was purified by flash chromatography, eluting with 25-50% ethyl acetate/petroleum spirit followed by 5-7% MeOH/DCM. *Purification method C*: The compound was purified by preparative reversed phase HPLC to afford the product as a bis-trifluoroacetate salt.

***Synthetic Methods***

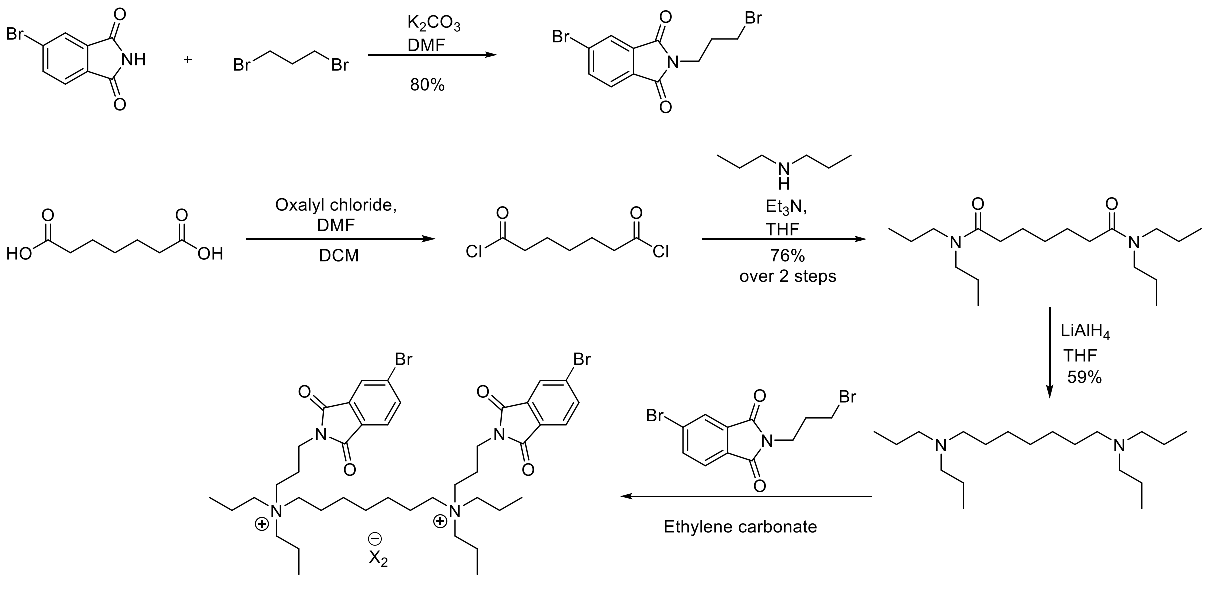


**Scheme 1:** Synthesis of 4P-C7/3-bromo-phth.

**5-Bromo-2-(3-bromopropyl)isoindoline-1,3-dione**

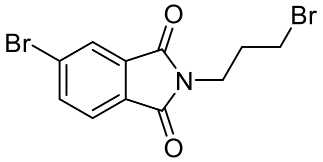

4-Bromophthalimide (500 mg, 2.21 mmol) and $K_2CO_3$ (336 mg, 2.43 mmol) were dissolved in 0.8 mL of DMF. 1,3-Dibromopropane (0.67 mL, 6.64 mmol) was then added and the solution stirred at 25 °C for 12 h. After completion of reaction, the solution was evaporated and EtOAc (20 mL) was added. The solution was then washed successively with $H_2O$, $NH_4Cl$ and brine. The organic layer was then dried with $MgSO_4$ and evaporated to dryness to obtain a white solid which was recrystallized in methanol to obtain the desired product as a white solid (66%, 511 mg, 1.5 mmol). $^1$H NMR (400 MHz, $CDCl_3$) δ 8.02 – 7.97 (m, 1H), 7.87 (dd, *J* = 7.9, 1.7 Hz, 1H), 7.74 – 7.69 (m, 1H), 3.84 (t, *J* = 6.9 Hz, 2H), 3.41 (t, *J* = 6.7 Hz, 2H), 2.26 (p, *J* = 6.7 Hz, 2H). $^{13}$C NMR (101 MHz, $CDCl_3$) δ 167.6, 167.0, 137.2, 133.8, 130.7, 129.2, 126.9, 124.9, 37.2, 31.6, 29.7. LC–MS *m/z* (relative intensity): $[M]^{1+}$ 347.6 (100%). HRMS (ESI–TOF) *m/z*: $[M]^{1+}$ Calcd for $C_{11}H_9Br_2NO_2$ 345.9073; found 345.9071.

***N*$^1$,*N*$^1$,*N*$^7$,*N*$^7$-tetrapropylheptanediamide**[1]

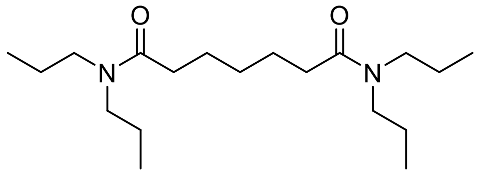


Title compound was synthesised using General procedure A using heptanedioic acid (1 g, 6.2 mmol) as a yellow oil (75%, 1.53 g). $^{1}$H NMR (400 MHz, $CDCl_3$) δ 3.31 – 3.22 (m, 4H), 3.22 – 3.12 (m, 4H), 2.47 – 2.19 (m, 4H), 1.74 – 1.46 (m, 12H), 1.44 – 1.31 (m, 2H), 0.89 (dt, *J* = 14.5, 7.4 Hz, 12H).

**$N^1$,$N^1$,$N^7$,$N^7$-tetrapropylheptane-1,7-diamine**[1]

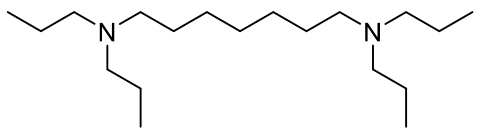


Title compound was synthesised using General procedure B from $N^1$,$N^1$,$N^7$,$N^7$-tetrapropylheptanediamide (100 mg, 0.306 mmol) as a colourless oil (49%, 45.2 mg). $^{1}$H NMR (400 MHz, $CDCl_3$) δ 2.45 – 2.27 (m, 12H), 1.49 – 1.43 (m, 12H), 1.25 – 1.35 (m, 6H), 0.86 (t, *J* = 7.4 Hz, 12H).

**$N^1$,$N^7$ -bis(3-(5-bromo-1,3-dioxoisoindolin-2-yl)propyl)- $N^1$,$N^1$,$N^7$,$N^7$-tetrapropylheptane-1,7-diaminium ditriflate** (4P-C7/3-bromo-phth)

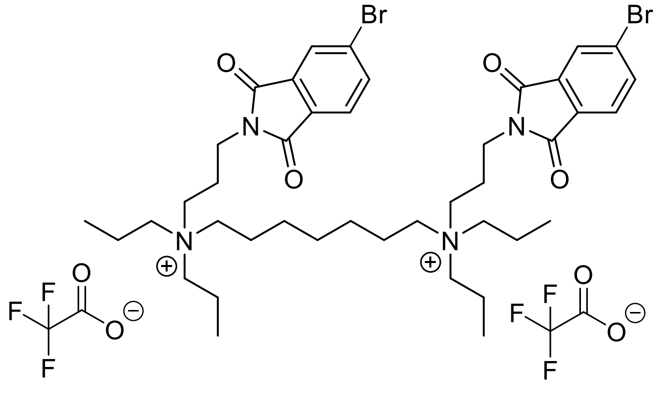


Title compound was synthesised using General procedure C from $N^1$,$N^1$,$N^7$,$N^7$-tetrapropylheptane-1,7-diamine (150 mg, 0.5 mmol). The crude product was successively purified using methods A, B, C and B to obtain the product as a clear oil (4.3%, 22.8 mg). $^{1}$H NMR (400 MHz, $CDCl_3$) δ 7.91 (d, *J* = 1.0 Hz, 2H), 7.84 (dd, *J* = 7.9, 1.5 Hz, 2H), 7.69 (d, *J* = 7.9 Hz, 2H), 3.78 (t, *J* = 6.5 Hz, 4H), 3.48 – 3.29 (m, 8H), 3.26 – 3.13 (m, 8H), 2.20 –2.07 (m, 4H), 1.73 – 1.60 (m, 12H), 1.44 – 1.31 (m, 6H), 0.97 (t, *J* = 7.1 Hz, 12H). $^{13}$C NMR (101 MHz, $CDCl_3$) δ 167.6 (2C), 167.1 (2C), 137.4 (2C), 133.5 (2C), 130.5 (2C), 129.3 (2C), 126.9

(2C), 125.0 (2C), 60.7 (4C), 59.5 (2C), 56.7 (2C), 35.3 (2C), 27.4, 25.4 (2C), 21.6 (2C), 21.4 (2C), 15.7 (4C), 10.8 (4C). $^{19}$F NMR (376 MHz, $CDCl_3$) δ -75.80. Purity (LC–MS, 254 nm): >94%. LC–MS *m/z* (relative intensity): $[M]^{2+}$ 415.9 (100%). HRMS (ESI–TOF) *m/z*: $[M]^{2+}$ Calcd for $C_{41}H_{60}Br_2N_4O_4^{2+}$ 416.1477; found 416.1496.

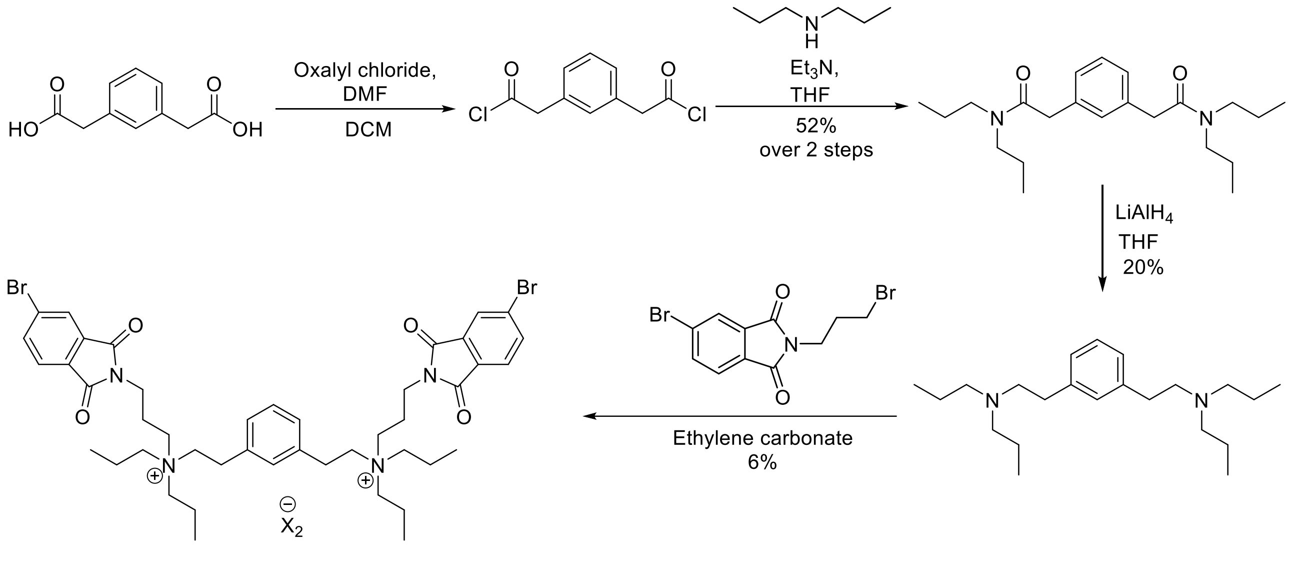


4B-aryl-C7/3-bromo-phth

**Scheme 2:** Synthesis of 4P-aryl-$C_7$/3-bromo-phth.

**2,2'-(1,3-phenylene)bis(*N*,*N*-dipropylacetamide)**

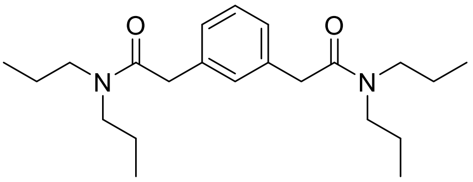

Title compound was synthesised using General procedure A using 1,3 phenylenediacetic acid (1 g, 5.1 mmol) as a yellow oil (52%, 955 mg). $^{1}$H NMR (400 MHz, $CDCl_3$) δ 7.28 – 7.21 (m, 2H), 7.15 – 7.11 (m, 2H), 3.67 (s, 4H), 3.24 (ddd, *J* = 42.7, 11.6, 6.8 Hz, 8H), 1.62 – 1.45 (m, 8H), 0.86 (td, *J* = 7.4, 1.1 Hz, 12H). $^{13}$C NMR (101 MHz, $CDCl_3$) δ 170.8 (2C), 136.0 (2C), 129.3, 129.0, 127.2 (2C), 50.1 (2C), 47.8 (2C), 40.9 (2C), 22.3 (2C), 21.0 (2C), 11.5 (2C), 11.3 (2C). LC–MS *m/z* (relative intensity): $[M + H]^{+}$ 361 (100%). HRMS (ESI–TOF) *m/z*: $[M]^{1+}$ Calcd for $C_{22}H_{36}N_2O_2$ 361.285; found 361.2856.

***N,N'*-(1,3-phenylenebis(ethane-2,1-diyl))bis(*N*-propylpropan-1-amine)**

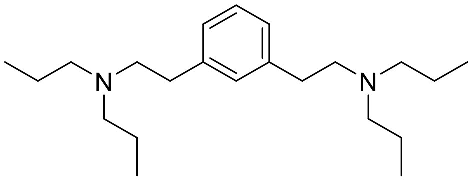

Title compound was synthesised using General procedure B from 2,2'-(1,3-phenylene)bis(*N,N*-dipropylacetamide) (500 mg, 1.39 mmol) as a yellow oil (20%, 94.6 mg). $^{1}$H NMR (400 MHz, $CDCl_3$) δ 7.21 – 7.15 (m, 1H), 7.04 – 6.98 (m, 3H), 2.81 – 2.62 (m, 8H), 2.57 – 2.38 (m, 8H), 1.60 – 1.39 (m, 8H), 0.89 (t, *J* = 7.4 Hz, 12H). $^{13}$C NMR (101 MHz, $CDCl_3$) δ 140.9 (2C), 129.3, 128.4, 126.4 (2C), 56.2 (6C), 33.5 (2C), 20.4 (4C), 12.1 (4C). LC–MS *m/z* (relative intensity): $[M + H]^+$ 333.1 (100%). HRMS (ESI–TOF) *m/z*: $[M]^{1+}$ Calcd for $C_{22}H_{40}N_2$ 333.3264; found 333.3268.

***N,N'*-(1,3-phenylenebis(ethane-2,1-diyl))bis(3-(5-bromo-1,3-dioxoisoindolin-2-yl)-*N,N*-dipropylpropan-1-aminium) ditriflate** (4P-aryl-$C_7$/3-bromo-phth)

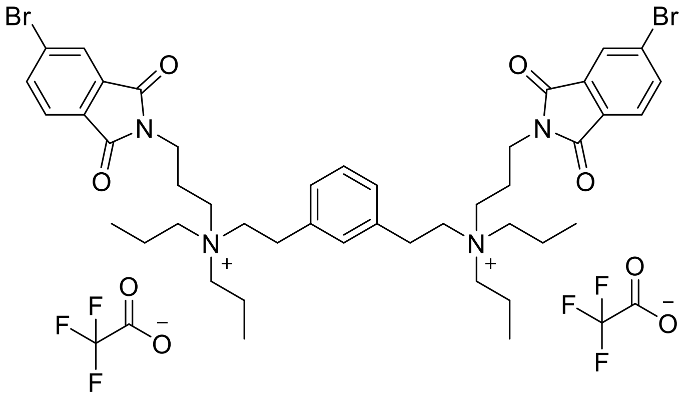

Title compound was synthesised using General procedure C from *N,N'*-(1,3-phenylenebis(ethane-2,1-diyl))bis(*N*-propylpropan-1-amine) (76 mg, 0.23 mmol). The reaction mixture was successively purified using method B and C to obtain the product as a colourless oil (6%, 15 mg). $^{1}$H NMR (400 MHz, $CDCl_3$) δ 7.91 ( br s, 2H), 7.84 (d, *J* = 7.7 Hz, 2H), 7.67 (d, *J* = 7.8 Hz, 2H), 7.25 – 7.14 (m, 2H), 7.13 – 7.03 (m, 2H), 3.82 – 3.68 (m, 4H), 3.41 (d, *J* = 28.7 Hz, 8H), 3.30 – 3.12 (m, 8H), 3.04 – 2.87 (m, 4H), 2.25 – 2.04 (m, 4H), 1.82 – 1.59 (m, 8H), 0.97 (t, *J* = 6.4 Hz, 12H). $^{13}$C NMR (101 MHz, $CDCl_3$) δ 167.5 (2C), 167.0 (2C), 137.5 (2C), 135.8 (2C), 133.5 (2C), 130.4 (2C), 130.0, 129.4 (2C), 129.1, 128.2 (2C), 126.9 (2C), 125.0 (2C), 61.1 (4C), 60.0 (2C), 57.0 (2C), 35.1 (2C), 28.2 (2C), 21.5 (2C), 15.7 (4C), 10.4 (4C).$^{19}$F NMR (376 MHz, $CDCl_3$) δ -75.78. Purity (LC–MS, 254 nm): >73%. LC–

MS *m/z* (relative intensity): $[M]^{2+}$ 432.9 (100%). HRMS (ESI–TOF) *m/z*: $[M]^{2+}$ Calcd for $C_{44}H_{58}Br_2N_4O_4^{2+}$ 433.1399; found 433.1414.

**$^{1}$H NMR Spectra for 4B-aryl-$C_7$/3-bromo-phth**

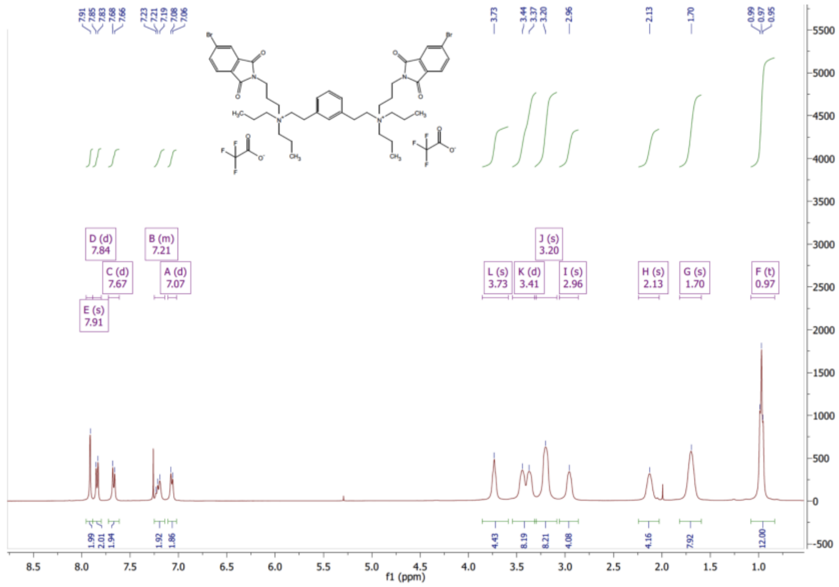


**$^{1}$H NMR Spectra for 4B-$C_7$/3-bromo-phth**

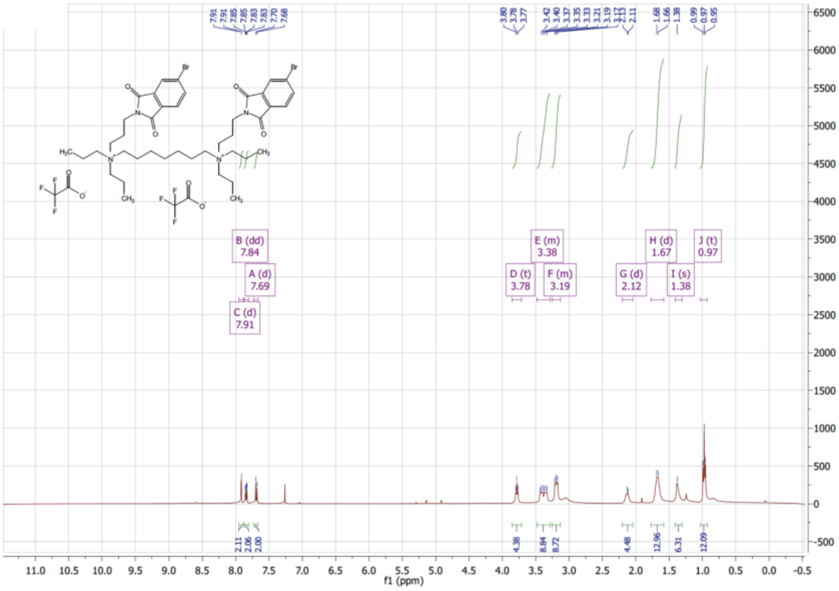
